# Mothers letting go: postnatal maternal investment shapes sex-specific social development in wild vervet monkeys

**DOI:** 10.64898/2026.02.16.705899

**Authors:** Josefien A. Tankink, Nokubonga Dlamini, Erica van de Waal

## Abstract

Sex differences in behaviour and life-history trajectories are widespread across species, yet the mechanisms through which mothers shape these differences remain poorly understood. Classic theories emphasize sex-biased allocation at birth or differential energetic investment, but how maternal effects might operate instead through postnatal investment remains understudied. Using long-term demographic and behavioural data from female-philopatric vervet monkeys (*Chlorocebus pygerythrus*), we examined how maternal age and dominance rank influence offspring sex ratios at birth, survival to adulthood, maternal investment, and offspring social integration in both sons and daughters. Maternal age, but not rank, influenced offspring sex ratios, with older females producing more daughters. Maternal rank was positively associated with daughters’ survival and social engagement, with estimated effects consistently stronger in daughters than in sons. While both sexes were highly vulnerable to maternal loss, post-hoc trends suggested a potentially steeper effect on sons. Sons received more maternal proximity (under some maternal conditions) and maternal grooming, whereas daughters seemed to gain earlier and greater engagement with other group members and appeared to derive indirect advantages from maternal rank through social exposure. Together, these findings indicate that maternal investment in this species differs in form rather than in magnitude, primarily through postnatal developmental pathways rather than biased allocation at birth. By demonstrating how maternal age and social status shape divergent early-life trajectories, our study highlights the role of early social environments in generating sex-specific life histories.

**Highlights:**

- Sex differences in life-history trajectories can arise through postnatal social development, not only sex allocation at birth;
- In wild female-philopatric vervets, maternal age (not rank) predicts offspring sex ratios, with older females producing more daughters;
- Maternal rank shows stronger associations with daughters’ survival and social engagement, while sons were potentially more dependent on maternal presence;
- Maternal investment differs in form rather than magnitude, consistent with role-specific developmental preparation.

## Introduction

Sex differences in behaviour, particularly in social strategies, are persistent among sexually reproducing animals (Kappeler, 2017). These differences in behaviour are largely driven by anisogamy, creating an initial asymmetry that leads to distinct evolutionary pathways towards the highest fitness (Kokko & Jennions, 2008; Schärer et al., 2012). While widespread, these differences in behaviour are not fixed, but are flexible and shaped by ecological constraints, social systems and the costs and benefits of parental care for each sex (Kappeler, 2017). In fact, mothers – the general prime caregiver of young in mammals (Clutton-Brock, 1991) – are known to adapt their behaviour, both towards their offspring as well as towards other group members, based on the offspring’s sex (Guinness et al., 1979; Hewison & Gaillard, 1999; Ishizuka & Inoue, 2023; Koskela et al., 2009; Kulik et al., 2016; Lonsdorf, 2017; Maestripieri, 2018; Murray et al., 2014; Robert et al., 2010). The mechanisms through which mothers shape these sex-specific trajectories can vary; maternal effects do not need to be expressed primarily through sex allocation at birth or energetic provisioning but may instead operate through postnatal social development and social shaping.

Sex-biased parental investment, both pre- and post-natal, in offspring is a long-standing research area in evolutionary biology (starting with Darwin, 1871). According to the parental investment theory (Trivers, 1972), the relative investment (i.e., any investment by a parent in an individual offspring that increases that offspring’s survival) in the sexes shapes patterns of competition and mate choice (also called “conventional sex roles”, Kokko & Jennions, 2008). An often discussed application of this framework is the Trivers-Willard hypothesis (TWH; Trivers & Willard, 1973), which predicts how mothers should adjust their investment both pre- and post-natal so that the sex ratio of their offspring is based on their own physical condition or social rank. In polygynous species, where male reproductive success is expected to vary more and is potentially more dependent on individual quality – while females usually reproduce regardless of condition, high-quality females should bias investment towards sons (Trivers & Willard, 1973). However, life-history analyses highlight that such predictions depend on maternal age, residual reproductive value, and the predictability of fitness returns, rather than condition alone (Leimar, 1996). In these models, mothers may favour the sex with more reliable or immediate reproductive payoffs when future reproductive opportunities decline, or when social and ecological constraints affect the expected benefits of producing sons versus daughters (Leimar, 1996). In species with sex-biased dispersal, maternal investment strategies can be further shaped by the social consequences of offspring residency, since offspring sex determines future exposure to kin competition and access to social allies. The “local resource competition” (LRC) hypothesis proposes that mothers may favour the dispersing sex when local competition is high, particularly for low-ranking females (Clark, 1978). In contrast, the “local resource enhancement” (LRE) hypothesis predicts that investment in the philopatric sex may be beneficial when offspring strengthen the matriline or provide social support, especially for high-ranking females (Emlen et al., 1986). These frameworks, while not mutually exclusive, emphasize that sex-biased maternal investment is a flexible strategy that integrates both internal state and external social structure to maximize lifetime fitness returns.

Maternal investment into their offspring can be expressed through a wide range of mechanisms operating across different developmental stages, both prenatally and postnatally. Mothers may adjust the sex ratio of their offspring before birth based on their own physical condition or social rank, to maximize fitness returns (Trivers & Willard, 1973). In tammar wallabies (*Notamacropus eugenii*), for example, mothers with higher “investment ability” were significantly more likely to give birth to sons (Robert et al., 2010). Maternal effects can also indirectly shape offspring phenotypes through prenatal hormonal exposure (Quinlivan et al. 1998; Hansen et al. 1999; Lesage et al. 2001, 2004; Seckl 2001; Walker et al. 2001), influencing later behavioural predispositions (Wallen & Hassett, 2009). After birth, mothers may further adjust their investment through the direct transfer of resources or physical effort to offspring based on their sex; producing richer or more milk for a specific sex (Hinde, 2009; Koskela et al., 2009), or directly adjusting their proximity and grooming behaviour towards offspring (Bentley-Condit, 2003; Fairbanks & McGuire, 1987; Kulik et al., 2016).

In primates in particular, these postnatal investment patterns are closely linked to the development of sex-specific social trajectories. Philopatric immature females form stronger bonds with maternal kin than males do (Amici et al., 2019; Lonsdorf, 2017; Maestripieri, 2018), while males often seek contact with other males or age-peers, potentially to form alliances and prepare for dispersal (Crockett & Pope, 1993; Lonsdorf, 2017; Maestripieri & Ross, 2004). Mothers may actively facilitate these divergent pathways by shaping offspring social exposure (Amici et al., 2019; Castellano-Navarro et al., 2023; Maestripieri, 2018), as shown in male-philopatric chimpanzees, where mothers of sons spend more time in groups containing adult males (Murray et al., 2014); potentially to prepare them for their future social environment (Lonsdorf, 2017). Likewise, mothers might encourage son’s dispersal in female-philopatric species by exhibiting higher rates of aggression towards sons (Kulik et al., 2016; Timme, 1995), while forming generally stronger bonds with their daughters (the philopatric sex; e.g., Ishizuka & Inoue, 2023; Kulik et al., 2016). These patterns suggest that maternal investment goes beyond energetic provisioning, but also prepares offspring with their future sex-specific roles.

Primates provide a strong study system for examining sex-biased maternal investment and its developmental consequences. Their slow life histories, extended periods of maternal care and complex social systems allow maternal effects to potentially accumulate over long developmental windows, making them useful for linking early experiences to adult social roles and potential fitness outcomes (Lonsdorf, 2017). Moreover, female philopatry in many primate species creates predictable sex differences in dispersal, kin competition and coalitionary behaviour, offering a natural context in which to study the predictions of TWH, LRC and LRE (Clark, 1978; Emlen et al., 1986; Trivers & Willard, 1973). Within this framework, long-term field studies are uniquely positioned to study variation in maternal age, rank, offspring survival and social integration. Vervet monkeys (*Chlorocebus pygerythrus*) provide an excellent model for examining sex-biased maternal investment within a female-philopatric primate social system. Females remain in their natal groups and form stable, matrilineal dominance hierarchies, whereas males disperse at maturity and must establish rank and social relationships in new groups (Borgeaud et al., 2016; Cheney & Seyfarth, 1990; Hemelrijk et al., 2020), forming dynamic multimale/multifemale groups. Maternal dominance rank in vervets has been found to predict the majority of conflicts of offspring (Horrocks & Hunte, 1983). Mating is polygynous, with little reproductive skew (Cheney et al., 1988; Minkner et al., 2018; Weingrill et al., 2011), and highly seasonal (Cheney et al., 1988). Females usually give birth to their first offspring when they are three years old, and then continuously one offspring per year, being a fast generation turnover for primates and enabling us to have a large sample size for this study. Mothers carry their infants for approximately the first three months, but infants gain slowly more independence during that period (Fairbanks & McGuire, 1987), while also receiving high interest and alloparental care from other group members, mainly females (Fruteau et al., 2011). The pronounced sex difference in dispersal creates predictable asymmetries in future social environments, kin competition, and fitness returns. Long-term data from well-habituated wild-living groups therefore allow energetic investment and social scaffolding to be evaluated together within a single natural system.

In this study, we use long-term behavioural and demographic data from a wild-living population of vervet monkeys to examine how maternal age and dominance rank shape sex-biased developmental trajectories. Specifically, we test whether (i) offspring sex ratios at birth vary with maternal age and dominance rank, as predicted by Trivers-Willard-type allocation models and local resource competition/enhancement frameworks (Côté & Festa-Bianchet, 2001; Maestripieri, 2002; Trivers & Willard, 1973); (ii) maternal rank, age and presence influences survival to adulthood for sons and daughters, consistent with sex-specific vulnerabilities and rank-mediated advantages (Horrocks & Hunte, 1983; Meikle & Vessey, 1988); (iii) maternal investment across development – considered through indirect (reproductive pacing) and direct investment (maternal proximity, grooming and coalitionary support towards the offspring) – varies by offspring sex and maternal characteristics (Maestripieri, 2018); and (iv) offspring social exposure and engagement with other group members align with these maternal investment patters, as expected if early experience is tuned to future sex-specific roles (Ishizuka & Inoue, 2023; Kulik et al., 2016; Lonsdorf, 2017; Murray et al., 2014). By integrating these components within a single population, we aim to clarify whether sex-biased maternal strategies in this female-philopatric primate are best understood as differential energetic investment, different social scaffolding, or a combination of both. While Trivers-Willard models (Trivers & Willard, 1973) emphasize fitness maximization through condition-dependent sex allocation, our study tests whether maternal effects operate primarily through sex-specific social preparation rather than differential energetic investment. Gaining a better understanding of how sexes develop, diverge and are treated by their mothers and other group members will help us shed light on the development of primate sociality and the evolution of sex roles, which remains a poorly understood aspect of (human and nonhuman) primate behavioural evolution (Maestripieri, 2018).

## Methods

Our aim was to examine (i) whether offspring sex ratios at birth varied with maternal age and dominance rank; (ii) how maternal rank and maternal presence influenced offspring survival to adulthood in sons and daughters; (iii) how maternal age and rank shaped patterns of maternal investment across offspring development, including reproductive pacing, spatial association, grooming, and coalitionary support; and (iv) whether sex differences in offspring social exposure and engagement with group members were consistent with these maternal investment patterns.

### Data collection

We studied wild vervet monkeys at the iNkawu Vervet Project in Mawana Game Reserve, South Africa. While the project started in 2010, reliable behavioural data on four groups could be collected from 2012 onwards. Behavioural data collection protocols changed in 2022, giving us 10 years of consistent, reliable behavioural data (2012-2022). Behavioural data needed to calculate maternal rank remained consistent over the data collection protocols, allowing us to calculate maternal rank from 2012 to 2025. Demographic data collection remained consistent over the years, giving us 15 years of demographic data (2010-2025). For an overview of which data was used for which question, see Table 1. Data were collected in four neighbouring, well-habituated groups (AK: mean group size 25.66 individuals, BD: mean group size 50.83 individuals, KB: mean group size 17.98 individuals, NH: mean group size 37.98 individuals), for which all individuals were individually identified. Observers started data collection after passing several quality control tests, such as interobserver reliability scoring – passing at least 80% of the kappa coefficient – with the on-site scientific or field manager, as well as identification tests of all individual monkeys in the groups. Data collection was conducted six days a week, with an average of 6.8 hours over 4.2 observation days per group, by on average 2.4 observers, since 2012 (start of the data period).

**Table 1:** overview of data used for each part of our analyses, including the time-period used and the number of offspring and mothers.

| Question | Time-period | n offspring; n mothers |
| --- | --- | --- |
| (i) Sex allocation at birth | 2012-2025 | 330; 98 |
| (ii) Offspring survival | 2012-2023 | 304; 94 |
| (iii) Maternal investment (IBI; proximity; grooming; support) | 2012-2022 | 222; 81 |
| (iv) Social exposure and engagement offspring | 2012-2022 | 222; 81 |

Births, deaths and group composition were monitored daily. For each infant, sex was recorded whenever possible. Because males disperse at approximately 4 years old and females start to reproduce at around 3-4 years of age, we restricted our survival analyses to survival up to 3 years of age (which we considered adulthood), while both sons and daughters still resided in their natal groups.

Mothers’ ages were known from long-term records or estimated from reproductive history at the beginning of the project. Maternal dominance rank was estimated using Elo-rating procedures based on agonistic interactions among adult females and were averaged per calendar year; given the stability of female hierarchies in this species (Borgeaud et al., 2017; Cheney & Seyfarth, 1990). Behavioural data on mother-offspring interactions and agonistic behaviour were collected through ad libitum and group scan sampling. For each mother-offspring pair and observation year, we calculated several binomial response variables: (i) proximity of mother to the infant, where each scan observation where the infant was recorded within 5 meters of the mother was recorded as a success (1), compared to each scan performed on the mother regardless of the proximity to her offspring (0); (ii) grooming investment into the infant by the mother where each grooming bout where the mother groomed the infant was recorded as a success, compared to each time the mother was recorded grooming any group member; (iii) maternal coalitionary support to the infant, where each conflict where the mother supported the infant was recorded as a success, compared to the total number of conflicts in which the infant was involved; (iv) the amount of offspring social engagement and exposure, where the number of grooming interactions or conflicts in which the infant was involved were recorded as a success, compared to the total number of grooming interactions or conflicts recorded in the group; the average distance of a mother to adult group members during the first year after an infant’s birth, calculated as a continuous numerical value and modelled using a Gamma distribution. The average distance of a mother to adult group members (offspring social exposure) was only considered in the first year of an offspring’s life as juvenile vervets gain independence rather quickly (Fairbanks & McGuire, 1987), and females give birth to new offspring almost every year (∼1.3 years, this study). An observational year started after the (estimated) date of birth of the infant. Most births in our population were recorded between October and January (97.2 % of births).

### Statistical analyses

All analyses were conducted in R (version 4.3.2) using the packages *glmmTMB*, *lme4*, *DHARMa*, and *emmeans* (Brooks et al., 2017; Hartig, 2024; Lenth & Piaskowski, 2017). For each birth, we scored infant sex and infant survival as a binary variable. Inter-birth intervals were calculated in years for each mother-year combination and used in the sex-ratio analyses as a proxy for maternal reproductive pace and potential costs of producing a given sex. For the behavioural datasets, offspring age was expressed in years, and behavioural measures were expressed per year of age (1-3). For the statistical analyses, infant age was mean-centred, and mother age, rank, number of females and inter-birth intervals were standardized and centred.

To test whether maternal rank, age or number of females present in the group during the previous mating season influenced the probability of producing a specific sex, we fitted a binomial generalized linear mixed model per infant birth. Our response variable, sex of the infant (0 for sons, 1 for daughters) was fitted with a logit-link using a binomial family. As fixed effects, we included maternal rank, maternal age, number of females present in the group during the previous mating season and the group identity. We included random intercepts for mother identity and offspring birth year to account for repeated measures within mothers and cohort-level differences among birth years. Maternal age and rank were weakly negatively correlated (r ≈ −0.15), and PCA did not reveal a dominant shared axis. VIF values were near 1, confirming negligible collinearity, so both variables were used as separate predictors of maternal quality.

To test for sex-specific effects of maternal rank, -age, and -death during childhood on offspring survival to age 3, we fitted another binomial generalized linear mixed model with binary survival as our response variable. Only infants born before 2023 were included in survival analysis, ensuring that survival outcomes to age 3 were known at time of analysis. Our fixed predictor variables consisted of the sex of the infant in an interaction with both the mother’s rank, the mother’s age at birth and a binary variable indicating whether the mother died during the infant’s childhood. Group identity was included in the model to account for any group differences. Again, mother identity as well as birth year of the infant were included as random effects.

To test whether the interbirth interval was affected by the previous birth, we fitted a similar binomial generalized linear mixed model with binary reproduction (reproduced next mating season yes/no) as our response variable. As fixed predictors we used the sex of the infant and maternal rank and age, as well as the group identity. Here again, mother identity as well as birth year of the infant were used as random effects.

For the six behavioural responses measuring maternal investment and offspring social engagement, we modelled the number of successes out of the number of trials using a beta-binomial generalized linear mixed model, to accommodate overdispersion in proportional data, or used the above-mentioned Gamma distribution for maternal distance to adult group members. These models included only the 222 offspring that survived for at least one year, as we wanted to avoid non-full years of data collection. Offspring of which the mother died during childhood were not considered for the mother investment models after the mother’s death (but were considered for the group engagement models). All six models shared the same fixed-effect structure, which included the sex of the offspring in an interaction with the age of the infant and the rank and age of the mother, except maternal proximity to other group members, which did not include offspring age (since only the first year of offspring were used). Group identity was again included to account for group differences. We included the offspring identity as a random effect to account for repeated measures (except for maternal proximity to adult group members) and allowed mother identity to vary in both intercept and offspring age slope (mother identity was included as a random effect without the offspring age slope for maternal proximity to adult group members). We allowed the dispersion parameter to vary across observation years, to account for different levels of observation effort, wherever the model allowed convergence (for an overview of models, see supplementary materials Table S1). We used a beta-binomial error structure, which accommodates extra-binomial variation and provided well-behaved residuals; in all final models, dispersion tests indicated either acceptable dispersion or (mild) underdispersion.

Residual diagnostics were conducted using *DHARMa* (Hartig, 2024), including simulation-based residual plots, tests for overdispersion and outlier tests, with bootstrap-based checks where appropriate. The model testing mother’s proximity to the infant showed underdispersion, all other models behaved well. We assessed the significance of fixed effects using Type II Wald χ² tests obtained via anova’s. For models with significant or marginally non-significant interactions, we used *emmeans* (Lenth & Piaskowski, 2017) to estimate sex-specific slopes and to test pairwise contrasts of trends between sons and daughters.

## Results

Out of 604 infants born between 2010-2025, 125 could not be reliably sexed before disappearance. Out of the 479 sexed infants, 388 survived for at least one year, meaning that approximately 64% of infants survived until one year of age. Approximately 44% of sexed infants were female. 308 infants survived to three years of age, giving a survival rate to adulthood of around 51%. Reproducing females had an average of 3.19 offspring in their recorded lifetime, with a minimum of 1 and a maximum of 10. The average interbirth interval was 1.34 years for mothers that had more than one offspring. About half of potentially reproductive females (females above the age of 3 that reproduced at least once in their lifetime) reproduced each year, with a minimum of 0% in 2024 and a maximum of 77.4% in 2011. These demographic summaries include all infants observed between 2010-2025. Analyses below use variable-specific subsets because (i) maternal rank is available from 2012 onward, (ii) survival analyses are restricted to cohorts with 3 year of follow-up, and (iii) proximity analyses use data collected through 2022 prior to protocol changes (see Methods).

### (i) Sex ratio at birth

Of the 330 sexed infants from 98 mothers for whom maternal rank could be reliably calculated, ∼45% were female. The probability of a birth being daughter increased significantly with maternal age (χ² = 4.796, *p* = 0.029; Figure S1). In contrast, maternal rank and the number of females present during the mating season showed no evidence of influencing offspring sex (rank: χ² = 0.018, *p* = 0.894; number of females: χ² = 0.202, *p* = 0.653). Sex-ratio at birth did not differ among groups (χ² = 0.628, *p* = 0.890).

### (ii) Offspring survival

The likelihood of survival to adulthood tended to increase with maternal rank (χ² = 7.693, p = 0.006). While the interaction between maternal rank and offspring sex was nonsignificant (χ² = 1.454, p = 0.228), post-hoc trends indicated that daughters’ survival increased significantly with maternal rank (slope = 0.646 ± 0.232 SE, p = 0.005), whereas sons showed no effect of maternal rank (slope = 0.286 ± 0.207 SE, p = 0.167; Figure 1a). The difference between these slopes were however not significant (slope = 0.360 ± 0.299 SE, p = 0.228). Maternal loss significantly decreased the likelihood to survive to adulthood as well (χ² = 19.984, *p* < 0.0001), with again no significant difference between sons and daughters (χ² = 0.181, *p* = 0.671). Post-hoc comparisons showed that indeed both sons and daughters were less likely to survive after maternal loss, although the effect appeared numerically stronger in sons (sons: slope = −1.51 ± 0.409 SE, *p* = 0.0002; daughters: slope = −1.26 ± 0.445 SE, *p* = 0.0046; Figure 1c), while not being statistically different from each other. Maternal age, offspring sex itself and group identity showed no significant effects on offspring survival (maternal age: χ² = 0.005, *p* = 0.946; infant sex: χ² = 0.004, *p* = 0.951; group: χ² = 5.005, *p* = 0.171; Figure 1b).

**Figure 1:**
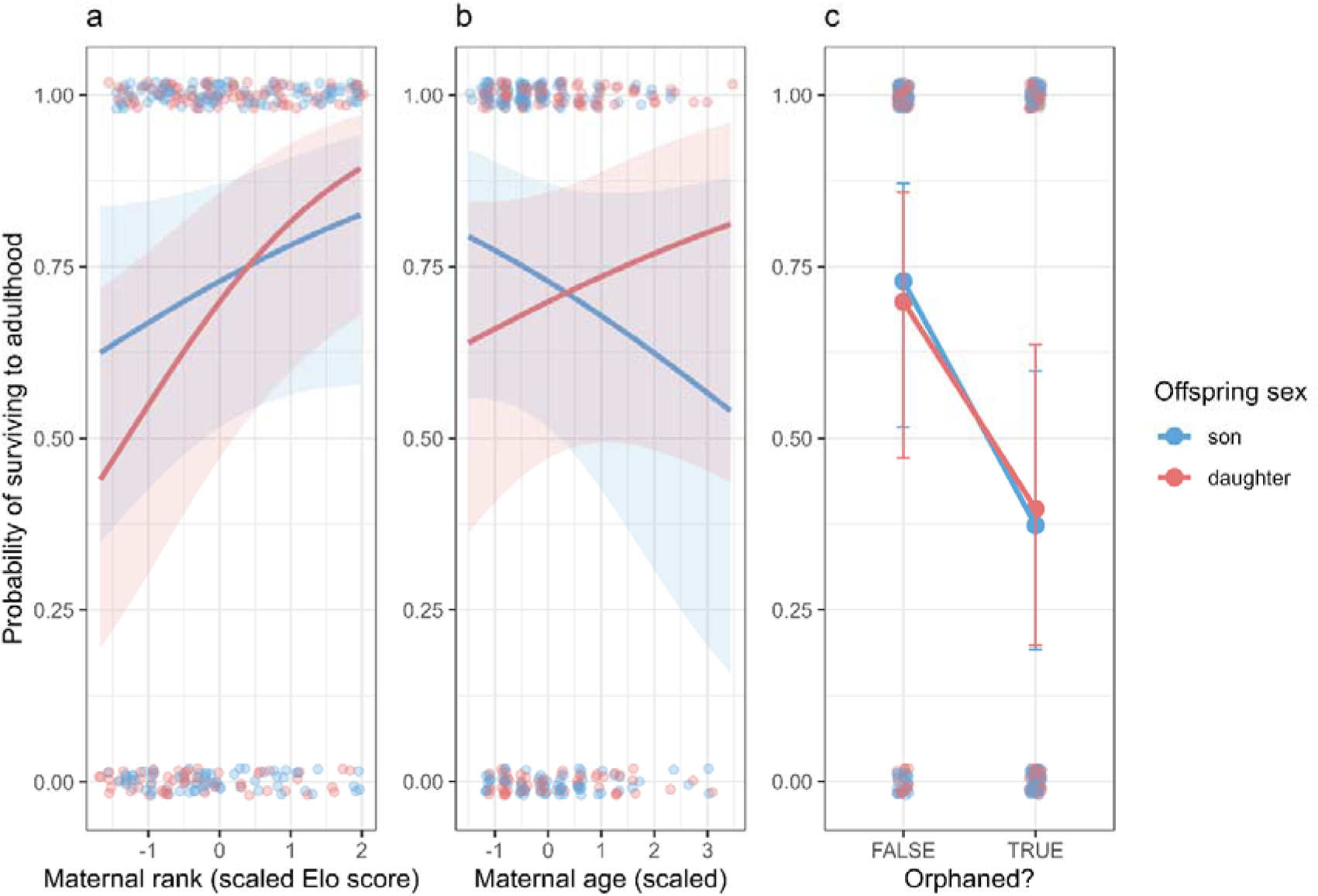
Predictors of offspring survival to adulthood (age 3) by offspring sex. Model-predicted probability of surviving to adulthood (age 3), shown separately for sons (blue) and daughters (red). Solid lines show predicted probabilities and shaded ribbons indicate 95% CIs; points show observed survival outcomes (0/1, jittered; raw data points). Points and error bars in (c) show predicted means ± 95% CI. (a) Maternal dominance rank (scaled Elo score): survival increased with maternal rank overall, and post-hoc trends indicated that daughters’ survival increased significantly with rank, whereas sons showed no clear rank effect. (b) Maternal age at birth (scaled): no evidence that maternal age predicted survival for either sex. (c) Maternal loss (Orphaned? FALSE/TRUE): offspring survival was strongly reduced by maternal death during childhood for both sexes; post-hoc trends suggested a steeper reduction in sons, but the sex difference in the orphaning effect was not significant.

### (iii-a) Reproductive pace

Mothers were marginally more likely to reproduce again the following breeding season after giving birth to daughters than to sons (χ² = 2.862, *p* = 0.091; Figure S2). Maternal rank, age and group identity did not affect the probability of a mother reproducing again the next breeding season (rank: χ² = 2.083, *p* =0.149; age: χ² = 1.392, *p* = 0.238; group: χ² = 6.050, *p* = 0.109; Figure S2).

### (iii-b) Maternal proximity to offspring

Maternal rank showed a significantly different effect on sons and daughters (χ² = 3.922, p = 0.048), where proximity to sons tended to increase with maternal rank (slope = 0.118 ± 0.069 SE, *p* = 0.086), whereas proximity to daughters tended to decrease, though not significantly (slope = −0.059 ± 0.075 SE, *p* = 0.429; Figure 2a). These sex-specific slopes differed significantly from one another (contrast = −0.177 ± 0.089 SE, *p* = 0.048). Maternal age had a significant overall effect on mother-offspring proximity (χ² = 8.897, *p* = 0.003). Although the interaction between offspring sex and maternal age did not reach conventional significance (χ² = 3.218, p = 0.073), post-hoc analyses showed that proximity to sons increased significantly with maternal age (slope = 0.224 ± 0.065 SE, *p* = 0.0005), whereas proximity to daughters showed no significant relationship with maternal age (slope = 0.075 ± 0.068 SE, *p* = 0.269; Figure 2b). The difference between these slopes was marginal (contrast = −0.148 ± 0.083 SE, *p* = 0.073). Mothers’ spatial association with offspring was not affected by offspring age (χ² = 0.033, *p* = 0.856; Figure 2c), and there was no evidence that proximity to the mother differed by offspring sex (χ² = 1.783, *p* = 0.182). Proximity to the mother varied among groups (χ² = 9.828, *p* = 0.020), with mothers in BD group generally maintaining lower proximity to their offspring than KB and NH group (BD – KB: slope = −0.500 ± 0.194 SE, *p* = 0.048; BD – NH: slope = −0.370 ± 0.143 SE, *p* = 0.047; Figure 6a). None of the other groups differed from each other. Random-effects estimates revealed substantial among-mother variation in baseline proximity and in offspring age-related slopes, as well as additional variance attributable to the offspring ID.

**Figure 2:**
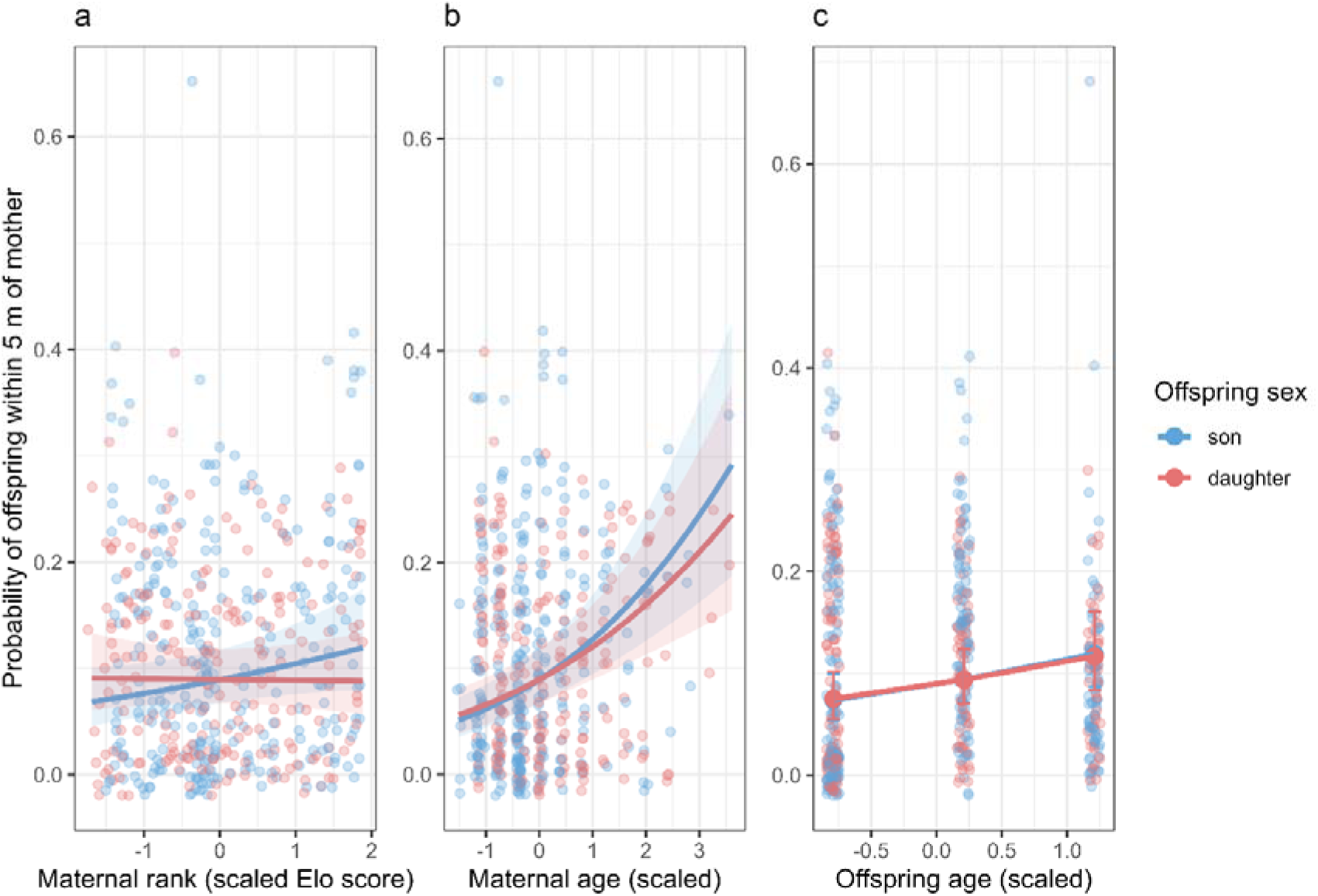
Maternal proximity to offspring across maternal rank, maternal age, and offspring age. Effects of maternal and offspring characteristics on the probability that offspring were observed within 5 m of their mother, shown separately for sons (blue) and daughters (red). Points represent individual offspring-year observations (jittered; raw data points); lines and shaded ribbons show model predictions ± 95% CI, and large points with error bars (c) indicate predicted means ± 95% CI. (a) Maternal dominance rank (scaled Elo score): rank showed a sex-specific effect (significant interaction), with proximity tending to increase with rank for sons but tending to decrease for daughters (slopes differed significantly). (b) Maternal age (scaled): proximity showed an overall positive effect of maternal age, driven by a significant increase for sons and no clear relationship for daughters (sex difference marginal). (c) Offspring age (scaled): proximity did not change with offspring age.

### (iii-c) Maternal grooming of offspring

Overall, sons received more maternal grooming than daughters (χ² = 4.163, *p* = 0.041; Figure 3). Maternal rank and maternal age had no significant main effects (rank: χ² = 0.104, *p* = 0.747; mother age: χ² = 1.359, *p* = 0.244; Figure 3a,b), and no interactions with sex were statistically significant (all interactions: *p* > 0.25). Mothers’ grooming investment in their offspring did not significantly change with offspring age (χ² = 2.526, *p* = 0.112), irrespective of offspring sex. While slopes for both infant sexes were not significantly different (χ² = 2.560, *p* = 0.110; slope contrasts = 0.112 ± 0.070 SE, *p* = 0.110), post-hoc trends suggested that maternal investment in offspring grooming increased with sons’ age (slope = 0.114 ± 0.051 SE, *p* = 0.026), while remaining flat for daughters (slope = 0.002 ± 0.058 SE, *p* = 0.973; Figure 3c). Group identity did not affect maternal grooming of offspring (χ² = 3.296, *p* = 0.348).

**Figure 3:**
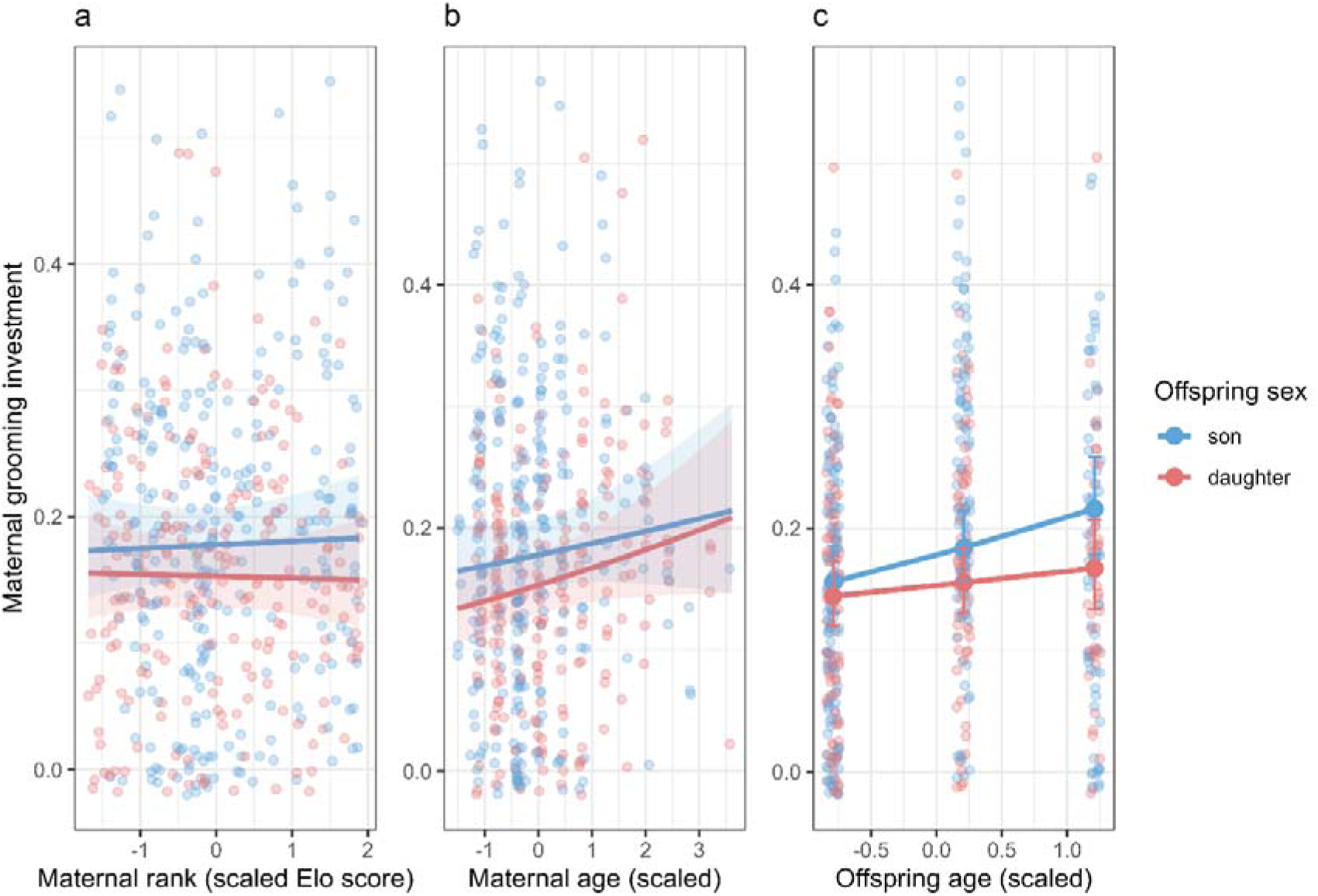
Maternal grooming investment toward offspring across maternal rank, maternal age, and offspring age. Maternal grooming investment (proportional grooming directed to the offspring) plotted against maternal and offspring predictors, shown separately for sons (blue) and daughters (red). Points represent individual offspring-year observations (jittered; raw data points); lines and shaded ribbons show model predictions ± 95% CI, and large points with error bars (c) indicate predicted means ± 95% CI. Across panels, mothers groomed sons more than daughters overall. (a) Maternal dominance rank (scaled Elo score): no evidence that rank predicted maternal grooming investment. (b) Maternal age (scaled): no evidence that maternal age predicted maternal grooming investment. (c) Offspring age (scaled): grooming investment did not show a significant age trend overall, but post-hoc trends suggested grooming increased with age for sons while remaining flat for daughters (sex-specific slope difference not significant). Across panels, mothers groomed sons more than daughters overall (main sex effect).

### (iii-d) Maternal coalitionary support

Mothers supported their offspring in conflicts in 3.2% of conflicts in which the offspring was involved. Mothers’ coalitionary support of their offspring was strongly affected by offspring age, maternal rank and maternal age (Figure S3). The probability that a mother supported her offspring decreased with offspring age (χ² = 16.539, *p* < 0.0001) and maternal age (χ² = 5.596, *p* = 0.018) but increased with mothers’ rank (χ² = 18.143, *p* < 0.0001), with no evidence that these patterns differed by offspring sex. The group identity and the offspring sex had no significant main effects (group: χ² = 1.901, *p* = 0.593; offspring sex: χ² = 0.042, *p* = 0.838). Interaction terms between sex and offspring age, maternal age or rank were not significant (all χ² ≤ 0.336, all *p* ≥ 0.562), indicating broadly similar patterns for both sons and daughters. However, post-hoc trends indicated that daughters were less likely to receive their mother’s support with increasing mother age (slope = −0.303 ± 0.138 SE, *p* = 0.028), but not sons (slope = −0.187 ± 0.155 SE, *p* = 0.228). The difference between these slopes was however not significant (estimate = 0.116 ± 0.200 SE, *p* = 0.562).

### (iv-a) Offspring social engagement

Female offspring were more engaged in both grooming interactions and conflicts than male offspring irrespective of age (grooming: χ² = 44.463, *p* < 0.0001; agonistic: χ² = 11.310, *p* = 0.0008; Figure 4). Maternal rank positively affected grooming engagement with group members of offspring, regardless of sex (χ² = 7.223, *p* = 0.007; Figure 4a), but not agonistic engagement (χ² = 0.576, *p* = 0.448; Figure 4d). Post-hoc trends revealed that while the interaction between offspring sex and maternal rank was not significant for grooming engagement (χ² = 0.688, *p* = 0.407), daughters showed a stronger positive effect (slope = 0.140 ± 0.053 SE, *p* = 0.009) than sons (slope = 0.093 ± 0.049 SE, *p* = 0.058). Maternal rank did not affect agonistic engagement of the sexes differently (χ² = 0.356, *p* = 0.551). Maternal age did not affect grooming (χ² = 1.429, *p* = 0.232), nor agonistic engagement of offspring (χ² = 0.004, *p* = 0.953; Figure 4b,e). The interaction between maternal age and offspring sex was not significant for both behaviours as well (grooming: χ² = 0.453, *p* = 0.501; conflicts: χ² = 0.134, *p* = 0.714). Post-hoc slopes confirmed that maternal age did not affect sons or daughters differently in both behaviours. The engagement in both grooming interactions and conflicts increased for both sexes with age (grooming: χ² = 402.082, *p* < 0.0001; conflicts: χ² = 369.964, *p* < 0.0001; Figure 4c,f) but showed a significant difference in slopes of grooming over offspring age by sex (χ² = 5.909, *p* = 0.015). Post-hoc trends revealed a stronger incline for daughters than for sons, while both sexes significantly inclined in their grooming with group members (sons: slope = 0.523 ± 0.039 SE, *p* < 0.0001; daughters: slope = 0.640 ± 0.037 SE, *p* < 0.0001). The interaction between offspring age and sex was not significant for agonistic engagement (χ² = 0.979, *p* = 0.322). Post-hoc slopes confirmed that age-related increases in conflict engagement were similar for sons and daughters.

**Figure 4:**
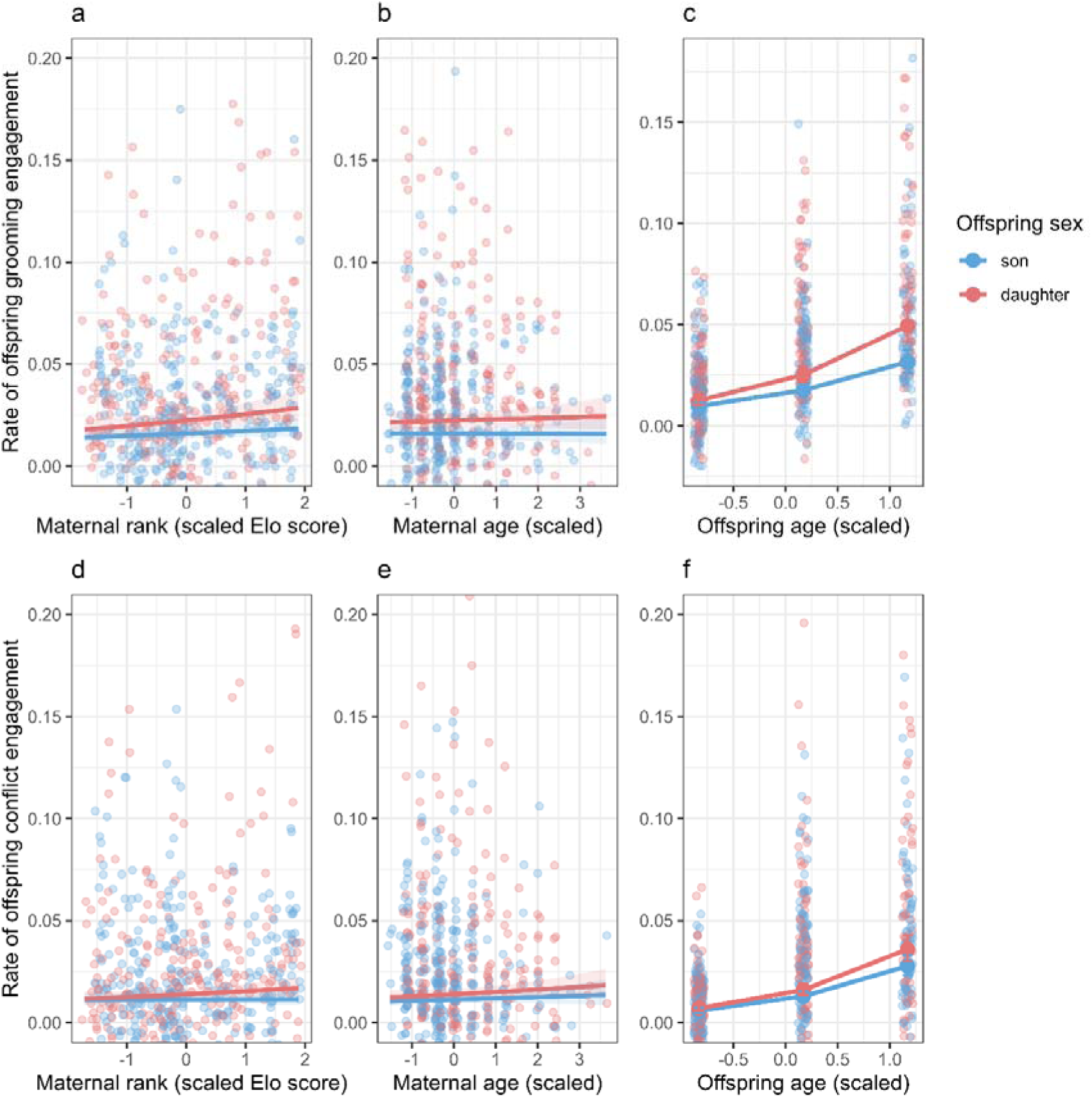
Offspring social engagement with group members across maternal rank, maternal age, and offspring age. Modelled offspring engagement rates in grooming (a-c, top row) and conflict (d-f, bottom row), shown separately for sons (blue) and daughters (red). Points show individual offspring-year observations (jittered; raw data points); lines and shaded ribbons show model predictions ± 95% CI, and large points with error bars (c, f) indicate predicted means ± 95% CI. Across panels, daughters were more engaged than sons. (a–c) Grooming engagement: (a) Maternal dominance rank (scaled Elo score): grooming engagement increased with maternal rank, with post-hoc trends indicating a clearer positive effect for daughters than sons, although the sex × rank interaction was not significant. (b) Maternal age (scaled): grooming engagement showed no relationship with maternal age. (c) Offspring age (scaled): grooming engagement increased strongly with offspring age in both sexes, with a steeper increase in daughters than sons (significant sex × age interaction). (d–f) Conflict engagement: (d) Maternal dominance rank (scaled Elo score): conflict engagement showed no relationship with maternal rank. (e) Maternal age (scaled): conflict engagement showed no relationship with maternal age. (f) Offspring age (scaled): conflict engagement increased strongly with offspring age for both sexes, with no sex difference in the age-related slope, although daughters were more engaged on average.

Group identity strongly affected offspring engagement with other group members (grooming: χ² = 82.377, *p* < 0.0001; conflicts: χ² = 106.943, *p* < 0.0001). Pairwise comparisons revealed pronounced differences among groups, with offspring in BD showing substantially lower engagement than those in AK, KB and NH in both behaviours. Offspring in KB showed higher engagement than those in AK and NH in both grooming and conflicts, whereas AK differed from KB but not from NH (Figure 6c,d).

### (iv-b) Exposure to other group members

Mothers differed consistently in how closely they positioned themselves relative to the nearest adult during the infant’s first year, with substantial among-mother variation in baseline spacing (random intercept variance = 0.055, SD = 0.235). There was no evidence that offspring sex predicted mean distance to the nearest adult (χ² = 0.488, *p* = 0.485). Maternal dominance rank, however, showed a marginal positive overall association with spacing (χ² = 3.277, *p* = 0.070). This effect did not differ by offspring sex (χ² = 0.287, *p* = 0.592). Post-hoc trend estimates indicated that distance tended to decrease with rank for mothers of sons (slope = −0.088 ± 0.050 SE, *p* = 0.080), whereas no significant relationship was detected for mothers of daughters (slope = −0.053 ± 0.053 SE, *p* = 0.317; Figure 5a). The difference between these slopes was however not significant (contrast = −0.035 ± 0.066 SE, *p* = 0.592). Maternal age had a strong overall increasing effect on spacing behaviour (χ² = 29.166, *p* < 0.001), with older mothers retaining lower proximity to adult neighbours. The effect of maternal age depended on offspring sex (χ² = 5.010, *p* = 0.025). Post-hoc analyses revealed that distance to the nearest adult increased strongly with maternal age for mothers of sons (slope = 0.290 ± 0.051 SE, *p* < 0.0001), whereas the increase with maternal age was weaker for mothers of daughters (slope = 0.140 ± 0.055 SE, *p* = 0.011; Figure 5b). These age-related slopes differed significantly between mothers of sons and daughters (contrast = 0.150 ± 0.067 SE, *p* = 0.025), indicating that the age-dependent increase in spacing was more pronounced when mothers were caring for sons. Mean distance to the nearest adult also varied among social groups (χ² = 11.338, *p* = 0.010), with BD-group-mothers having smaller distance to their nearest adult neighbour than KB- and (marginally) NH-group-mothers (BD – KB: slope = −0.502 ± 0.169 SE, *p* = 0.016; BD – NH: slope = −0.232 ± 0.099 SE, *p* = 0.089; Figure 6b).

**Figure 5:**
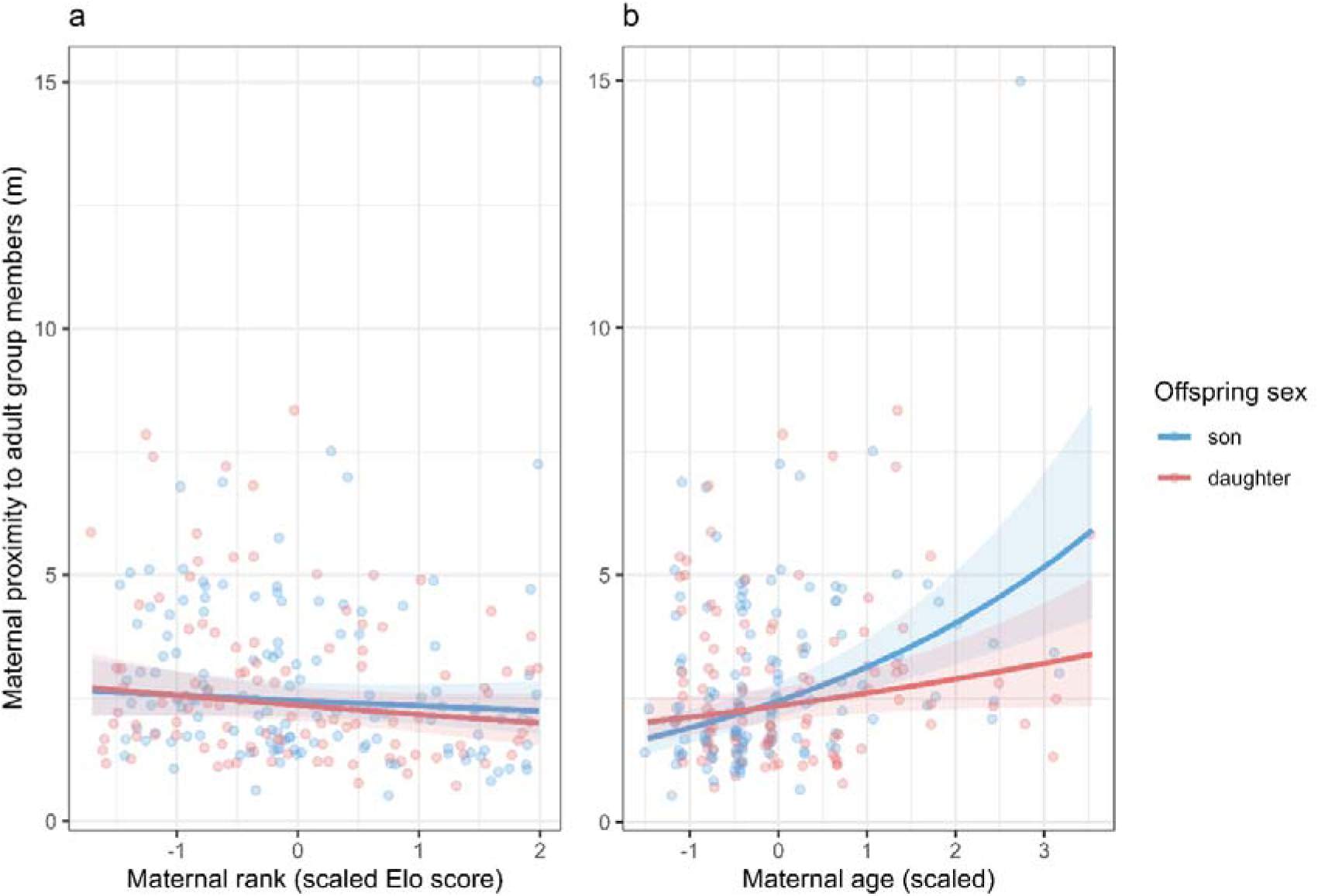
Maternal effects on proximity to adult group members during the offspring’s first year. Maternal proximity to adult group members (m; distance to the nearest adult neighbour) plotted as a function of maternal characteristics, shown separately for mothers of sons (blue) and daughters (red). Semi-transparent points show individual observations (jittered; raw data points). Lines show model predictions with shaded 95% confidence intervals. (a) Maternal rank (scaled Elo score): distance to the nearest adult showed a marginal overall association with rank, with post-hoc trends suggesting slightly smaller distances at higher rank for mothers of sons and no clear relationship for mothers of daughters; the rank × offspring sex interaction was not significant. (b) Maternal age (scaled): distance to the nearest adult increased strongly with maternal age, and this effect was sex-dependent (significant interaction), with a steeper age-related increase for mothers of sons than for mothers of daughters.

**Figure 6:**
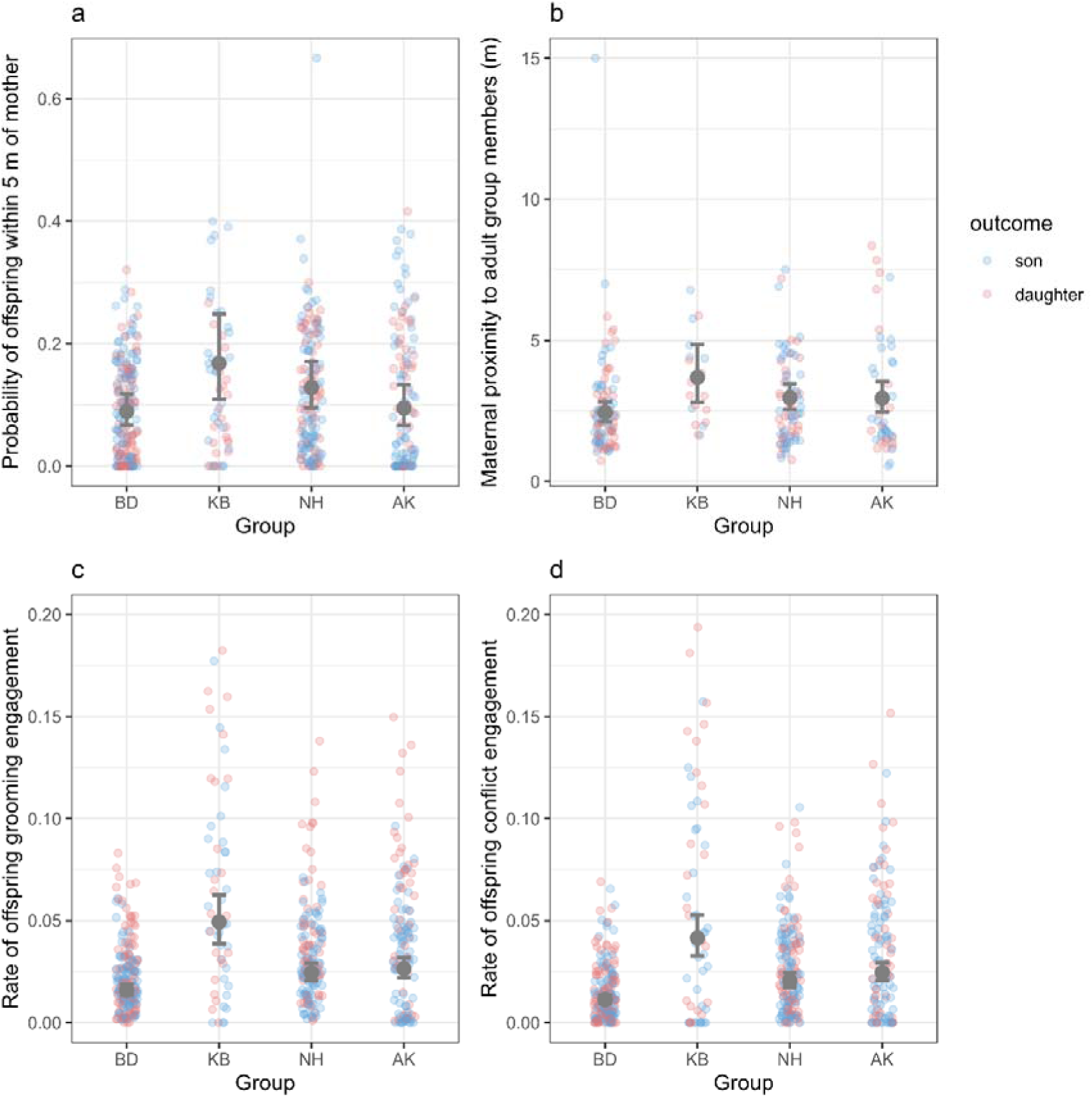
Group differences in mother–offspring proximity, maternal proximity to adult group members, and offspring social engagement. Group-level variation across the four study groups (BD, KB, NH, AK). Semi-transparent points show individual offspring-year observations (raw data; jittered; blue = sons, red = daughters). Large grey points with error bars show predicted group means ± 95% CI. (a) Probability that the offspring was observed within 5 m of its mother; mother-offspring proximity differed among groups (overall group effect), with mothers in BD generally showing lower proximity to their offspring than KB and NH. (b) Maternal proximity to adult group members (m; distance to the nearest adult neighbour during the offspring’s first year); spacing to the nearest adult differed among groups (overall group effect), with BD mothers maintaining smaller distances (closer proximity) to adults than KB (and marginally NH). (c) Offspring grooming engagement rate (proportion of group grooming interactions involving the focal offspring); offspring in BD showed lower grooming engagement than offspring in the other groups (overall group effect), and daughters were more engaged in grooming than sons. (d) Offspring conflict engagement rate (proportion of group agonistic interactions involving the focal offspring); offspring in BD showed lower conflict engagement than other groups (overall group effect), and daughters were more engaged in conflicts than sons.

## Discussion

Sex differences in behaviour and life-history trajectories are widespread across animals, yet the mechanisms through which mothers shape these differences remain poorly understood. Our long-term data suggest clear sex differences in how vervet offspring develop, interact with their mothers, and survive to adulthood. Together, these findings indicate that maternal investment in this system differs in form rather than in magnitude, and that maternal effects are expressed primarily through postnatal developmental pathways that shape offspring social exposure and engagement, rather than through biased allocation at birth alone.

We examined how maternal characteristics (age and dominance rank) shape sex allocation, offspring survival, maternal investment, and offspring social exposure, and how these processes differ between sons and daughters in a female-philopatric primate. Specifically, we examined (i) whether offspring sex ratios at birth varied with maternal age and dominance rank; (ii) how maternal rank and maternal presence influenced offspring survival to adulthood in sons and daughters; (iii) how maternal age and rank shaped patterns of maternal investment across offspring development, including reproductive pacing, spatial association, grooming, and coalitionary support; and (iv) whether sex differences in offspring social exposure and engagement with group members were consistent with these maternal investment patterns.

Maternal effects on offspring operated in distinctive ways in our population. Our results indicate that sons mainly benefit from the direct physical presence of their mother, reflected in the prolonged proximity and increased maternal grooming, while being potentially particularly vulnerable to maternal loss. Daughters, in contrast, showed patterns consistent with benefiting indirectly from their mother’s rank, and tended to engage earlier and more extensively with other group members. Although formal infant sex and maternal rank interactions were not consistently significant across models, the estimated rank-related effects were repeatedly larger in daughters than in sons. These patterns are consistent with the idea that maternal behaviour can function as role-specific developmental preparation of both sexes – i.e., social scaffolding – rather than reflecting only energetic bias.

Our result that older mothers were more likely to produce daughters, but not higher-ranking mothers, contrasts findings in cercopithecine primates (Maestripieri, 2002), but is in line with work on ungulates and other primates, where maternal age was a stronger predictor of sex allocation at birth than social status (Côté & Festa-Bianchet, 2001; Lonsdorf, 2017). These age-related biases towards producing daughters are consistent with models predicting increased investment in the sex with the more predictable fitness outcome, when future reproductive opportunities decline (Leimar, 1996); as well as with indications that daughters represent a “safer” investment for older females in female-philopatric primates (Brown, 2001). Although local resource competition and enhancement models (Clark, 1978; Emlen et al., 1986) are often discussed in the context of sex allocation, our findings suggest that in our population, these processes are expressed primarily through postnatal developmental pathways, rather than through biased sex ratios at birth.

More broadly, the pattern that daughters appear to gain greater developmental advantages from maternal rank is consistent with evidence from vervets and other primates, where maternal rank is directly transmitted to daughters and strongly determines their competitive and reproductive success (Horrocks & Hunte, 1983; Maestripieri, 2018; Meikle & Vessey, 1988). In our data, rank-related effects on survival and social engagement were directionally stronger in daughters, even when statistical tests of interactions did not always detect clear sex differences. This convergence across outcomes aligns with the local resource enhancement theory (Emlen et al., 1986), although expressed developmentally (postnatally) rather than through sex allocation (prenatally). At the same time, sons being more dependent on the presence of their mother is mirrored in findings in macaques and other primates, where male offspring were found to be more sensitive to maternal loss and early-life adversity (Meikle & Vessey, 1988; Patterson et al., 2024). Other evidence of mothers potentially actively facilitating social integration and independence in the philopatric sex comes from chimpanzees, where mothers of sons were found to spend more time with males in the first six months of their son’s life (Murray et al., 2014). While we did not find equally strong sex differences in all behavioural domains, the reduced proximity to other group members of older females with sons was absent when these females had daughters, indicating similar patterns. Since vervets reproduce offspring nearly annually and offspring remain semi-dependent on their mother for about three years (mirrored by the effect of maternal loss on survival of offspring to three years), it can be expected that these results would be less pronounced in this species compared to more slower-reproducing species. Nevertheless, the indication that daughters in our population engaged more in social interactions with other group members, whereas mothers maintained somewhat stronger direct grooming investment in sons, point in the same direction. Such early engagement may prepare daughters for lifelong residence within the group’s female hierarchy, whereas sons, who will disperse, may gain less from early integration into the maternal social and rank structure.

These patterns overall do not indicate reduced maternal care towards daughters per se, but rather suggest differences in how mothers allocate investment across sexes. Maternal age and rank appear to modulate whether investment is expressed through direct protection and care (in sons) or through indirect benefits mediated by social status and network integrations (in daughters). This aligns with Lonsdorf’s (2017) framework, which emphasizes that maternal effects on offspring fitness often arise through opportunities for social positioning rather than through differences in caregiving effort.

Notably, our results differ from several studies in other female-philopatric primate species that show stronger mother-daughter bonds and higher maternal investment in daughters (e.g., Ishizuka & Inoue, 2023; Kulik et al., 2016). Instead, vervet mothers in our population maintained stronger spatial and grooming relationships with sons. This suggests that sex-biased maternal strategies can be flexible and may depend on species-specific social dynamics, levels of aggression, or the degree to which maternal presence provides immediate protective benefits. Interbirth intervals were marginally shorter after the birth of a daughter, indicating lower reproductive cost. Whether this potential lower reproductive cost comes from a reduced maternal investment, or whether reduced maternal investment follows as daughters are less costly, remains unclear. However, it can be said that male vervets have a potentially higher reproductive output than females, since reproductive skew seems to be limited in this species (Minkner et al., 2018). In a seasonal system where females are constrained in annual reproductive output and males can achieve highly variable reproductive success, maternal investment may be shaped less by energetic ‘cost’ and more by the differential survival risks and social trajectories of sons versus daughters.

Some contrasting group differences were found in our proximity data as well: while mothers remained generally closer to other adult group members in BD compared to KB and NH, they remained further from their offspring in this group. Such patterns are consistent with previously documented group differences in social dynamics in this population, which persists despite broadly similar ecological conditions and demographic composition. Long-term data from the same study site indicate that groups differ in their overall levels of affiliation, likely reflecting group social styles rather than short-term demographic effects (Kerjean et al., 2024). Experimental work on co-feeding tolerance further supports the idea that groups can differ in how closely individuals tolerate proximity to others, particularly in contexts involving mothers and infants (Opreni et al., 2025). Since offspring engagement was lower in BD than in all other groups, mothers might feel safer to allow infants to range further while remaining near other adults. Although group size could contribute to these group differences, the patterns do not align with what we would expect if group size were the primary driver. Specifically, given that BD is consistently the largest group and KB the smallest, a group-size explanation would predict a consistent ordering across outcomes (BD > NH > AK > KB). However, the observed group differences do not follow this ordering across measures. This suggests that the group differences observed here likely reflect background variation in group-level social structure, rather than group size per se. Together, these findings suggest that the group differences observed here likely reflect background variation in group-level social structure, rather than maternal strategies or group sizes.

This study leverages long-term observational data, but several limitations should be considered. Because demographic, rank, and behavioural measures were not available uniformly across the full project, analyses rely on variable-specific time windows and partially overlapping samples, which complicates cross-outcome comparisons. Measures of maternal investment (proximity, grooming, coalitionary support) capture important components of care but do not include energetic investment or physiological measures, and coalitionary support in particular is rare, which limits power to detect sex-specific effects. Behavioural measures of investment are conditioned on observation opportunities and, particularly for rare behaviours such as coalitionary support, may have limited power to detect sex-specific moderation. Finally, although restricting behavioural analyses to 2012-2022 minimizes impacts of protocol changes, variation in observation effort and year- or group-level ecological conditions may still contribute to patterns observed in this single population. As in any observational study, unmeasured environmental variation and individual condition could confound associations with maternal rank and age, and interaction effects may be underpowered even in a long-term dataset.

Together, our findings indicate that mothers prepare sons and daughters for different social futures. Sons rely heavily on maternal proximity, grooming and protection, making them potentially extra vulnerable to maternal loss, whereas daughters develop social independence earlier and potentially capitalize on inherited rank and broader social connections. These divergent pathways arise not from strong sex-biases in maternal care, but from consistent differences in how maternal age, rank and behaviour translate into developmental opportunities for sons and daughters. By jointly examining sex allocation at birth, offspring survival, maternal behaviour and social engagement in multiple groups but within the same population, our results provide an integrated view of how maternal age, dominance rank and offspring sex interact to shape early life trajectories in a female-philopatric primate. Our study highlights how subtle developmental differences accumulate into the sex-specific life histories characteristic of many mammals and emphasizes the importance of early social environments in shaping the evolution of sex roles.

## Acknowledgements

We thank the onsite managers, Albert Driescher, Arend van Blerk, Michael Henshall, Siboniso Thela, Zonke Mbutho and all the field assistants, Master students, PhD students and postdocs who collected the data over the study period. We are grateful to the van der Walt family for giving us the permission to conduct the study on their land. This project was funded by the Swiss National Science Foundation (P300P3_151187, 31003A_159587, PP00P3_170624, PP00P3_198913 and CRSII-222818) along with Branco Weiss Fellowship–Society in Science, the grant ‘ProFemmes’ of the Faculty of Biology and Medicine, University of Lausanne and by the European Research Council under the European Union’s Horizon 2020 research and innovation programme for the ERC ‘KNOWLEDGE MOVES’ starting grant (grant agreement no. 949379) that also supported J.T., N.D. and E.v.d.W. during the time of analysing and writing.

For the purpose of Open Access, a CC BY public copyright license is applied to any Author Accepted Manuscript (AAM) version arising from this submission.

## AI declaration

AI was used during preparation of this manuscript solely to improve readability of the text and assist in refining coding. Any AI-generated material was meticulously reviewed, edited and verified by the authors. All ideas, interpretations and conclusions are from the authors. The authors take full responsibility for the accuracy and originality of all content.

## Author Contributions

J.A.T and E.vdW. conceived of the presented idea. N.D. oversaw continuous data collection. J.A.T. conceptualized data analysis and carried out data cleaning and analytic calculations. E.vdW. helped with the interpretation of the results. J.A.T. wrote the drafts of the manuscript that were edited and approved by all authors.

## Data availability and open access

all relevant data and code are available on OSF: https://osf.io/pekdj/overview?view_only=dcd93054cb4a4744b6c810565b3e6e67.

## Ethics statement

Data collection adhered to the ASAB Guidelines (Behaviour, 2018) and was purely observational. All individuals observed in this study were habituated to human presence and there were no interactions between humans and study subjects during the study. No permit is required for observational research on this species conducted on private land. Nevertheless, Ezemvelo KZN Wildlife and the van der Walt family, the owners of reserve where the study was conducted, approved the study and granted permission.

## Appendix

**Table S1:** Overview of statistical models used in this study.

| Topic | Response variable | Error distribution /link | Fixed effects (predictors) | Random effects | Notes / data subset |
| --- | --- | --- | --- | --- | --- |
| (i) Sex allocation at birth | Offspring sex at birth (daughter vs son) | Binomial / logit | Maternal rank + Maternal age + Number of adult females in group during the mating season + Group identity | Mother identity (random intercept) + Birth year (random intercept) | Includes only births with known offspring sex and maternal rank available. |
| (ii) Offspring survival | Survival to adulthood (age 3) (yes/no) | Binomial / logit | Offspring sex × (Maternal rank + Maternal age + Maternal loss during childhood) + Group identity | Mother identity (random intercept) + Birth year (random intercept) | Restricted to cohorts where survival outcome to age 3 was known (births before 2023). |
| (iii-a) Reproductive pace | Reproduction in the following breeding season (yes/no; proxy for short interbirth interval) | Binomial / logit | Offspring sex + Maternal rank + Maternal age + Group identity | Mother identity (random intercept) + Birth year (random intercept) | Response coded as whether the mother reproduced the next season (binary). |
| (iii-b) Maternal investment: proximity to offspring | Maternal proximity to offspring (proportion of scans where offspring was within 5 m of mother) | Beta-binomial / logit | Offspring sex × (Offspring age + Maternal rank + Maternal age) + Group identity | Mother identity (random intercept + random slope of offspring age) + Offspring identity (random intercept) | Behavioural data 2012–2022; only offspring surviving ≥1 year; models include repeated annual measures (offspring age 1–3). Dispersion modelled as a function of observation year. |
| (iii-c)<br>Maternal investment: grooming of offspring | Maternal grooming investment (proportion of mother's grooming directed to offspring) | Beta-binomial / logit | Offspring sex × (Offspring age + Maternal rank + Maternal age) + Group identity | Mother identity (random intercept + random slope of offspring age) + Offspring identity (random intercept) | Behavioural data 2012–2022; only offspring surviving ≥1 year; dispersion modelled as a function of observation year. |
| (iii-d)<br>Maternal investment: coalitionary support | Maternal coalitionary support (proportion of offspring conflicts in which mother supported offspring) | Beta-binomial / logit | Offspring sex × (Offspring age + Maternal rank + Maternal age) + Group identity | Mother identity (random intercept + random slope of offspring age) + Offspring identity (random intercept) | Behavioural data 2012–2022; trials restricted to offspring conflicts (trials > 0). Constant dispersion (no observation-year dispersion term). |
| (iv-a.1)<br>Offspring social engagement: grooming | Offspring grooming engagement (proportion of group grooming interactions involving the focal offspring) | Beta-binomial / logit | Offspring sex × (Offspring age + Maternal rank + Maternal age) + Group identity | Mother identity (random intercept + random slope of offspring age) + Offspring identity (random intercept) | Behavioural data 2012–2022; only offspring surviving ≥1 year; dispersion modelled as a function of observation year. |
| (iv-a.2)<br>Offspring social engagement: agonistic interactions | Offspring conflict engagement (proportion of group agonistic interactions involving the focal offspring) | Beta-binomial / logit | Offspring sex × (Offspring age + Maternal rank + Maternal age) + Group identity | Mother identity (random intercept + random slope of offspring age) + Offspring identity (random intercept) | Behavioural data 2012–2022; trials restricted to years with recorded conflicts (trials > 0); dispersion modelled as a function of observation year. |
| (iv-b)<br>Offspring<br>social<br>exposure:<br>maternal<br>proximity to<br>adult group<br>members | Maternal<br>proximity to adult<br>group members<br>(mean distance to<br>nearest adult<br>neighbour during<br>offspring's first<br>year; m) | Gamma /<br>log | Offspring sex ×<br>(Maternal age +<br>Maternal rank) +<br>Group identity | Mother identity<br>(random<br>intercept) | First-year measure<br>only (no offspring-age<br>term). Dispersion<br>modelled as a<br>function of calendar<br>year. |

**Table S2:**
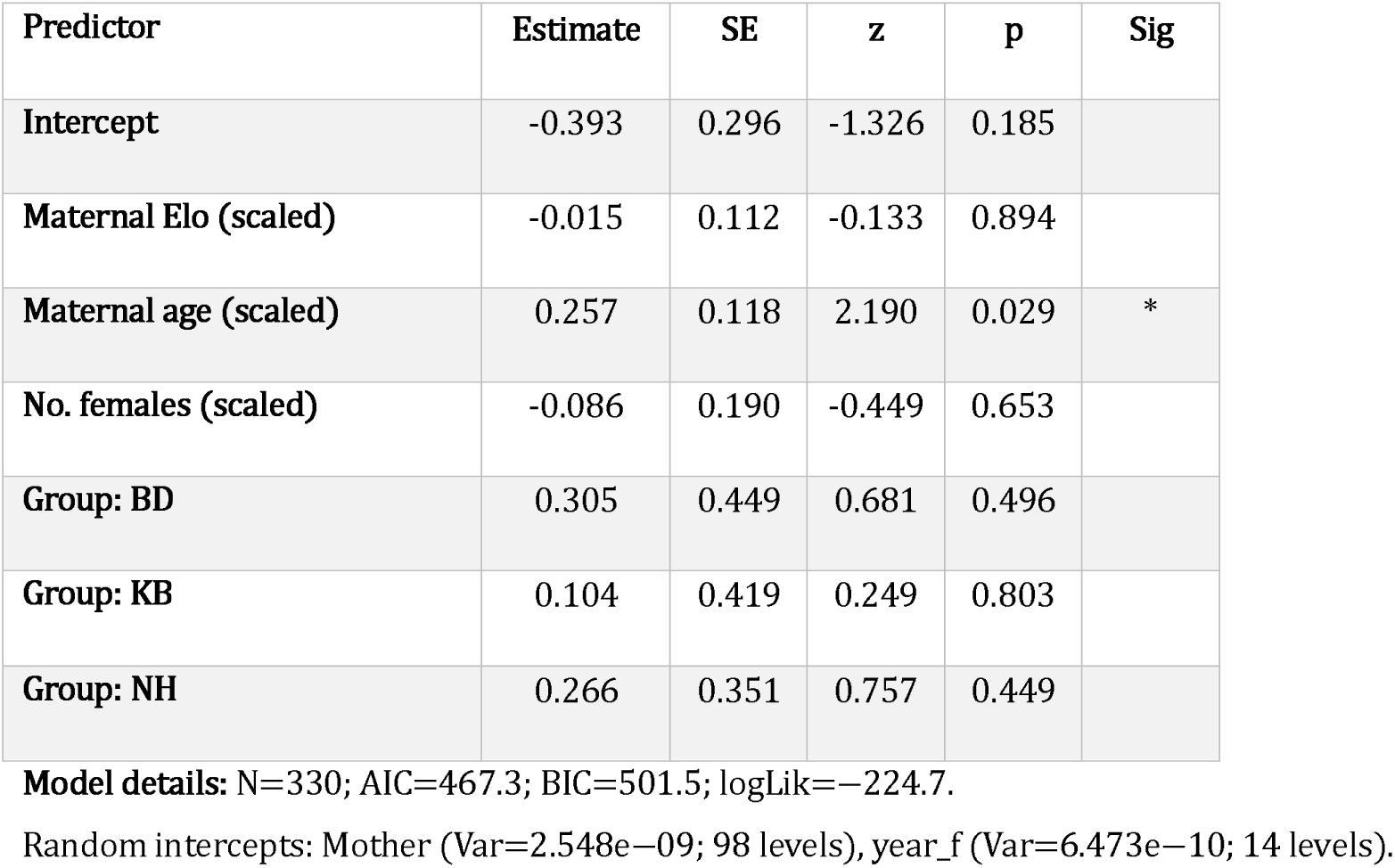
Birth sex ratio model (binomial GLMM; response = daughter).

| Predictor | Estimate | SE | z | p | Sig |
| --- | --- | --- | --- | --- | --- |
| Intercept | -0.393 | 0.296 | -1.326 | 0.185 |  |
| Maternal Elo (scaled) | -0.015 | 0.112 | -0.133 | 0.894 |  |
| Maternal age (scaled) | 0.257 | 0.118 | 2.190 | 0.029 | * |
| No. females (scaled) | -0.086 | 0.190 | -0.449 | 0.653 |  |
| Group: BD | 0.305 | 0.449 | 0.681 | 0.496 |  |
| Group: KB | 0.104 | 0.419 | 0.249 | 0.803 |  |
| Group: NH | 0.266 | 0.351 | 0.757 | 0.449 |  |

**Model details:** N=330; AIC=467.3; BIC=501.5; logLik=-224.7.
Random intercepts: Mother (Var=2.548e-09; 98 levels), year\_f (Var=6.473e-10; 14 levels).

**Table S3:**
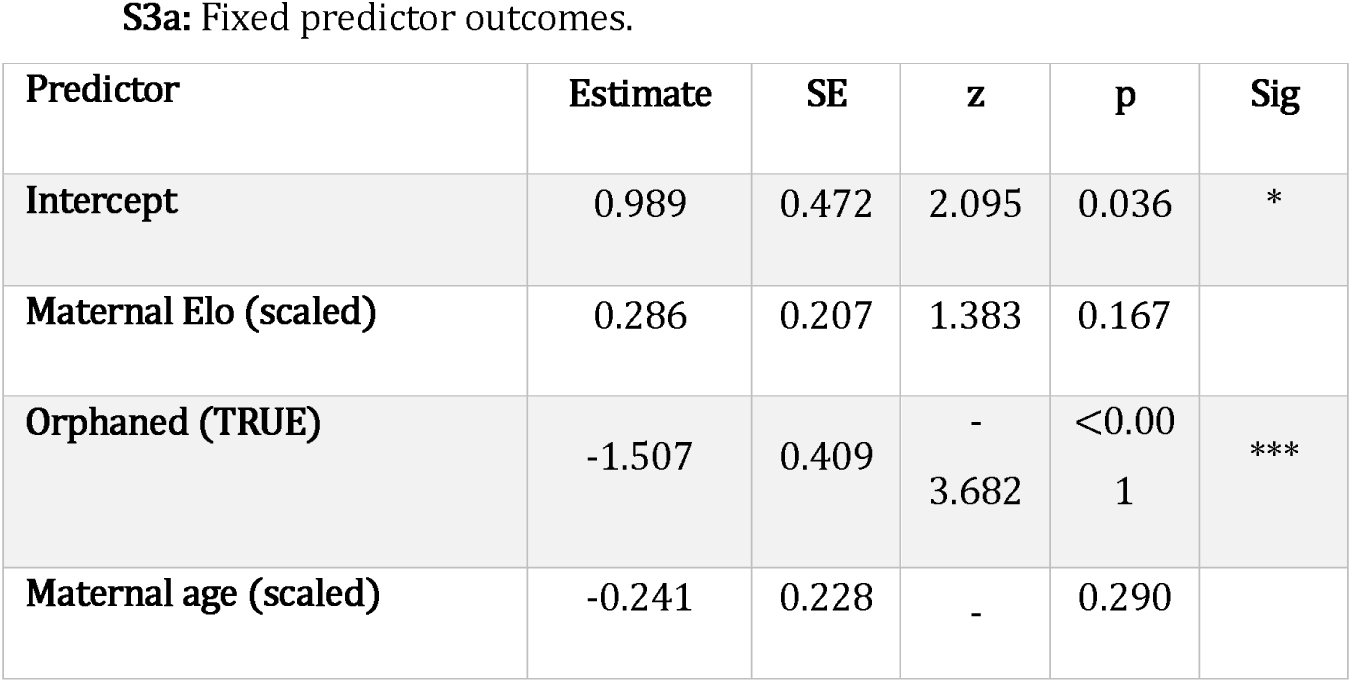
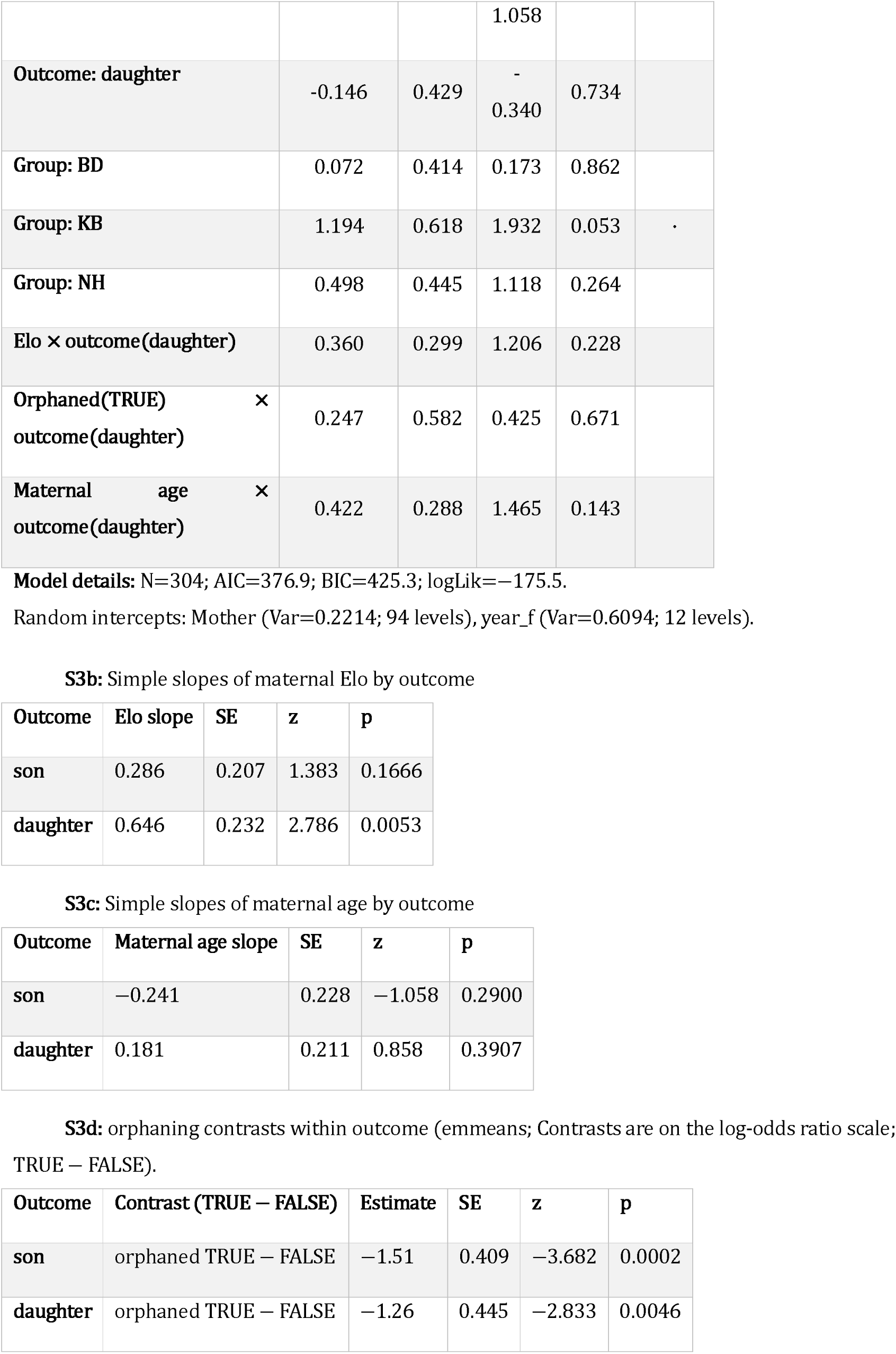
Survival model (binomial GLMM; response = survived_to_adult).

**Table S4:**
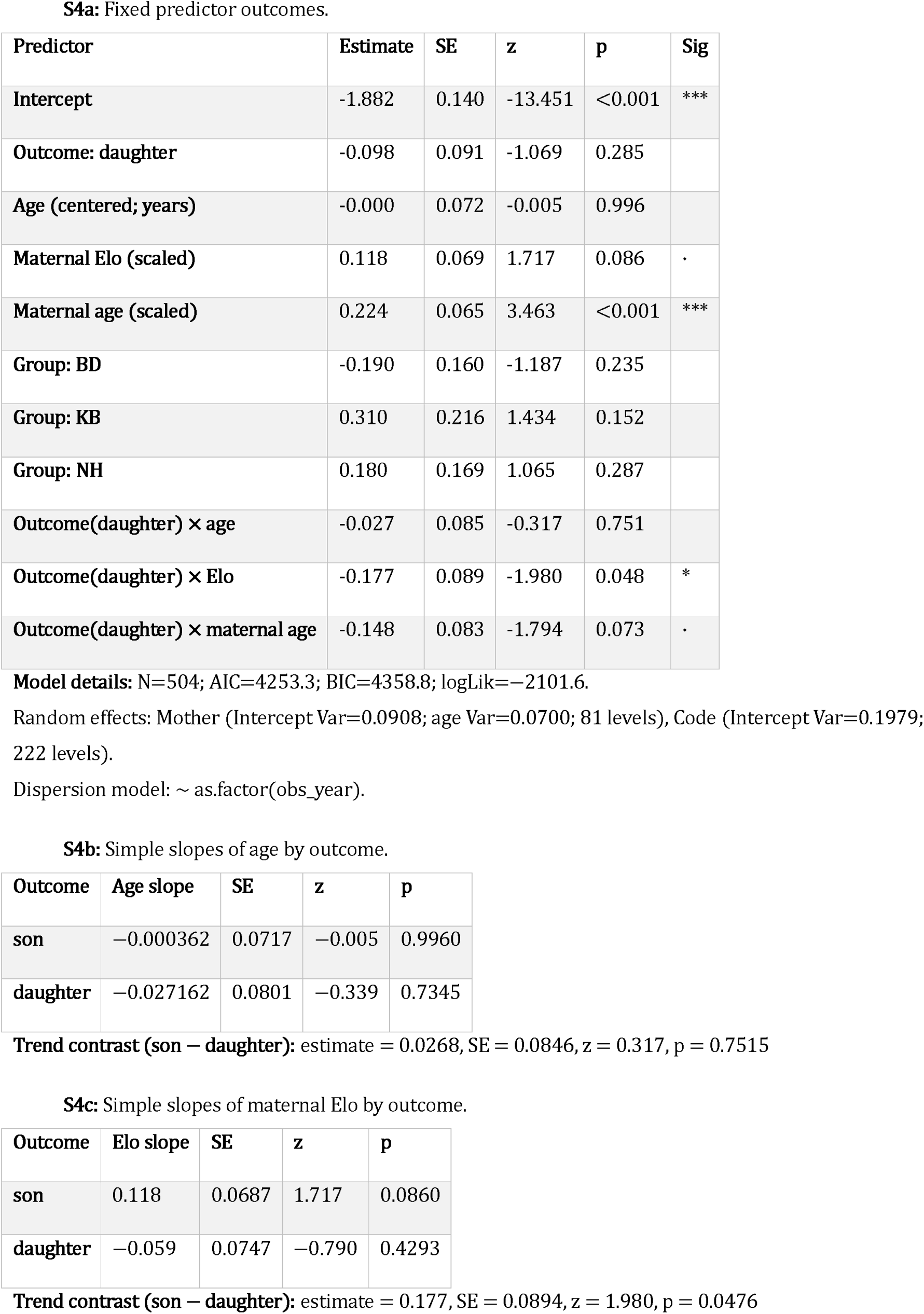
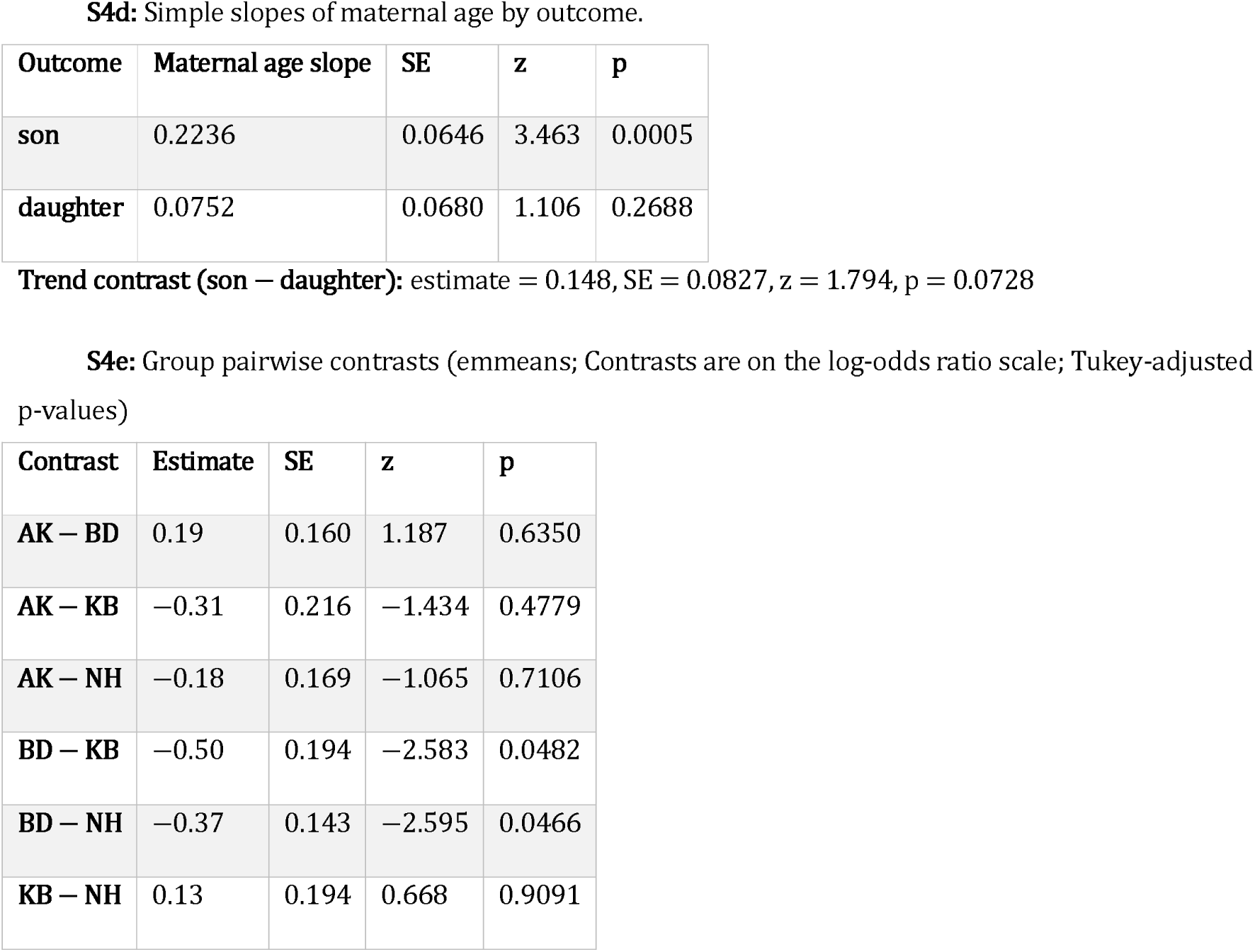
Proximity model (betabinomial GLMM; response = cbind(successes, trials−successes)).

**Table S5:**
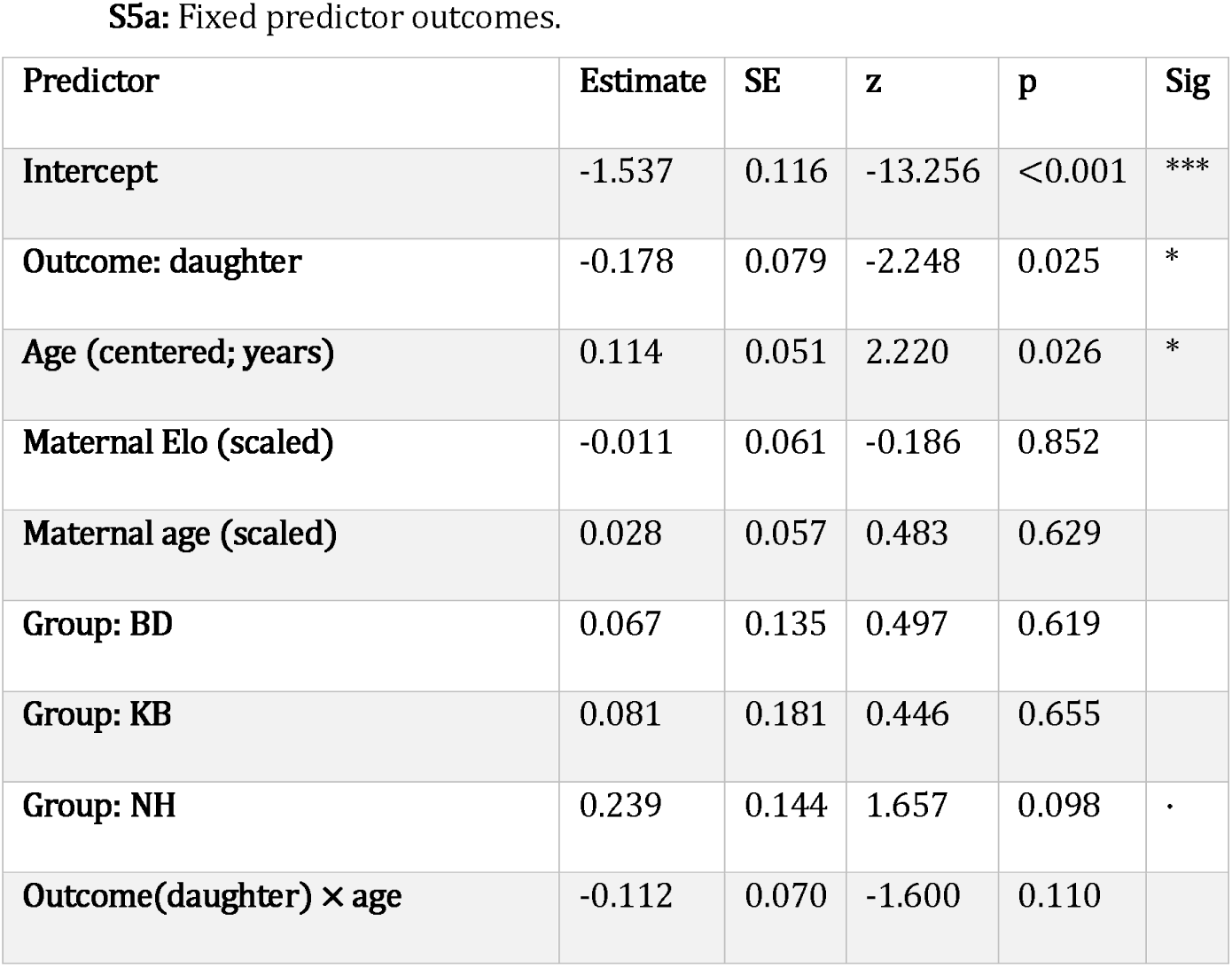
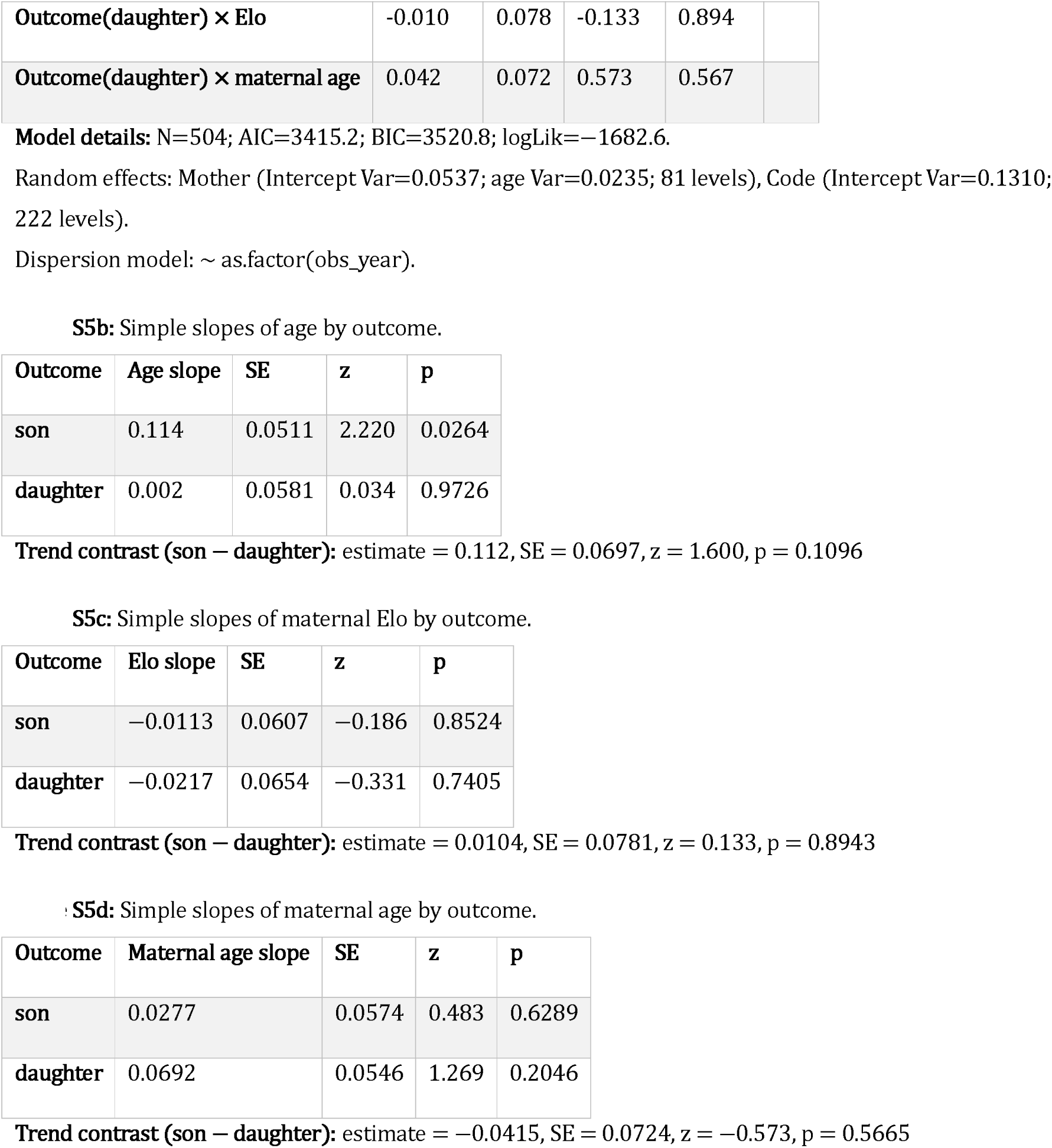
Maternal grooming model (betabinomial GLMM; response = cbind(successes, trials−successes)).

**Table S6:**
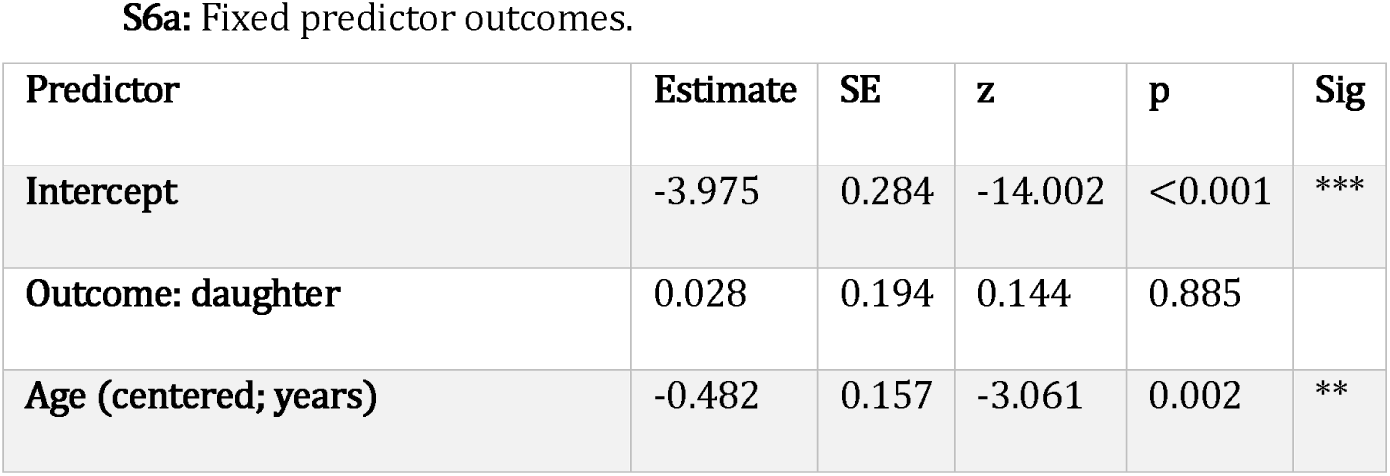
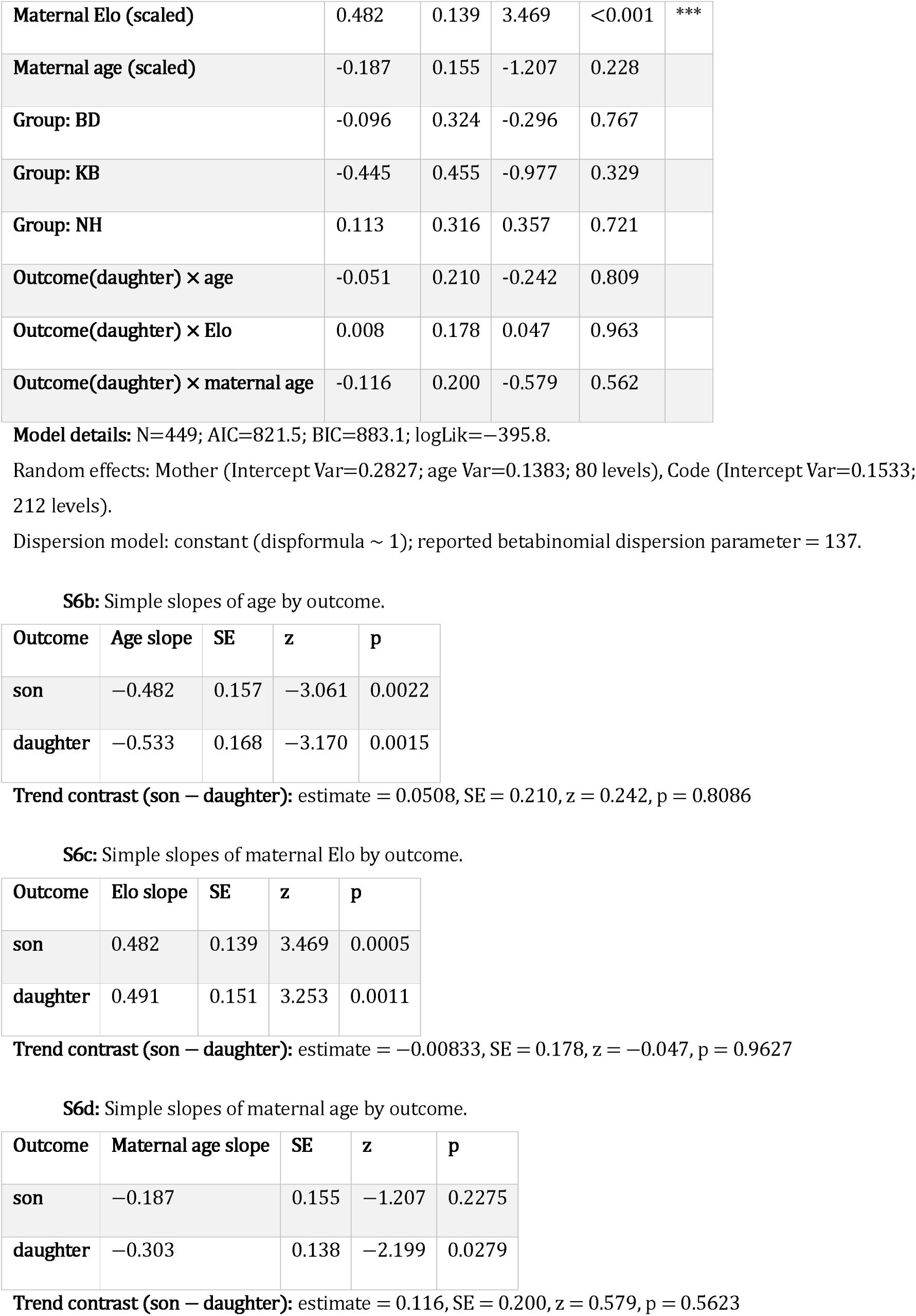
Coalitionary support model (betabinomial GLMM; ; response = cbind(successes, trials−successes; trials ≠ 0). Table S6a: Fixed predictor outcomes.

**Table S7:**
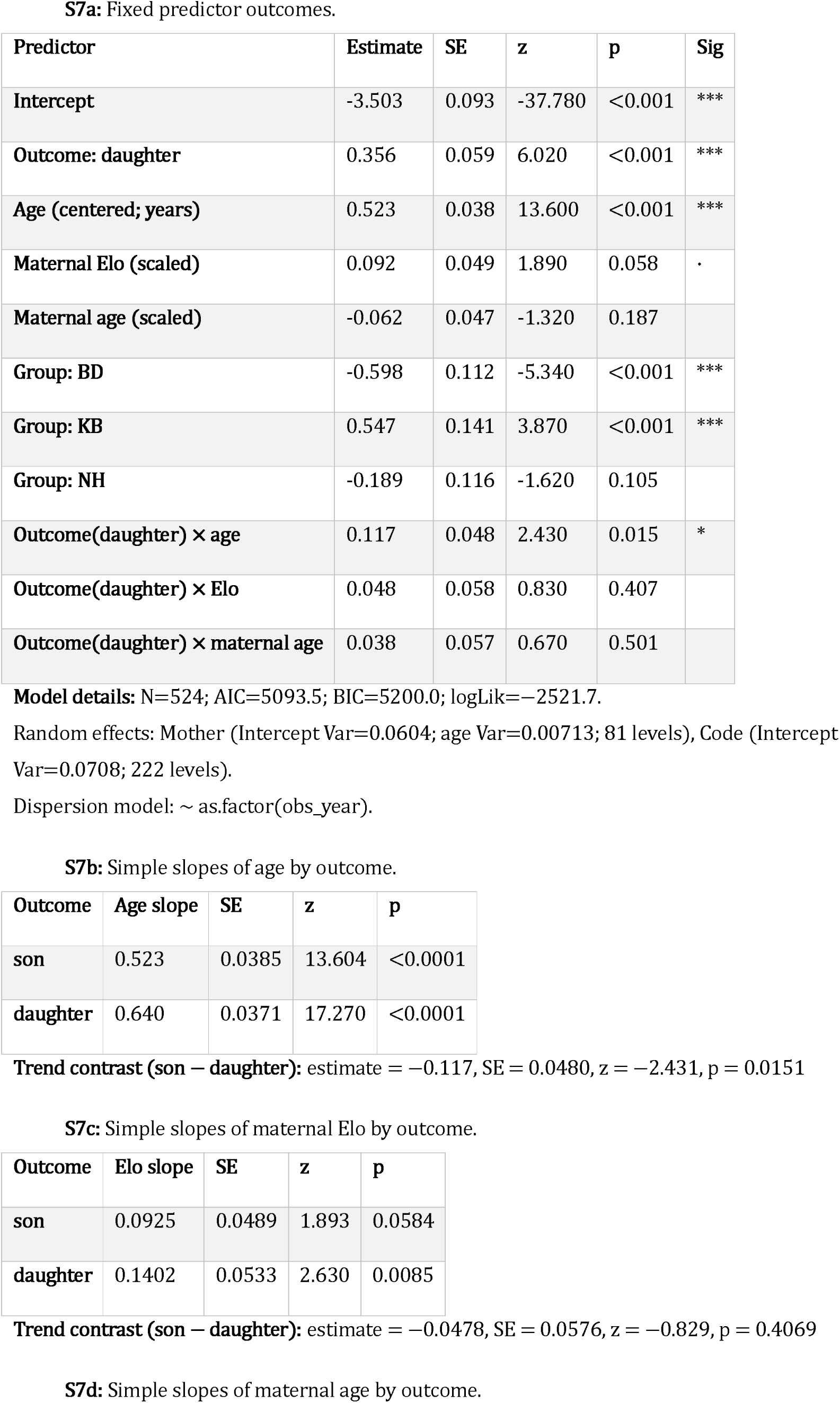
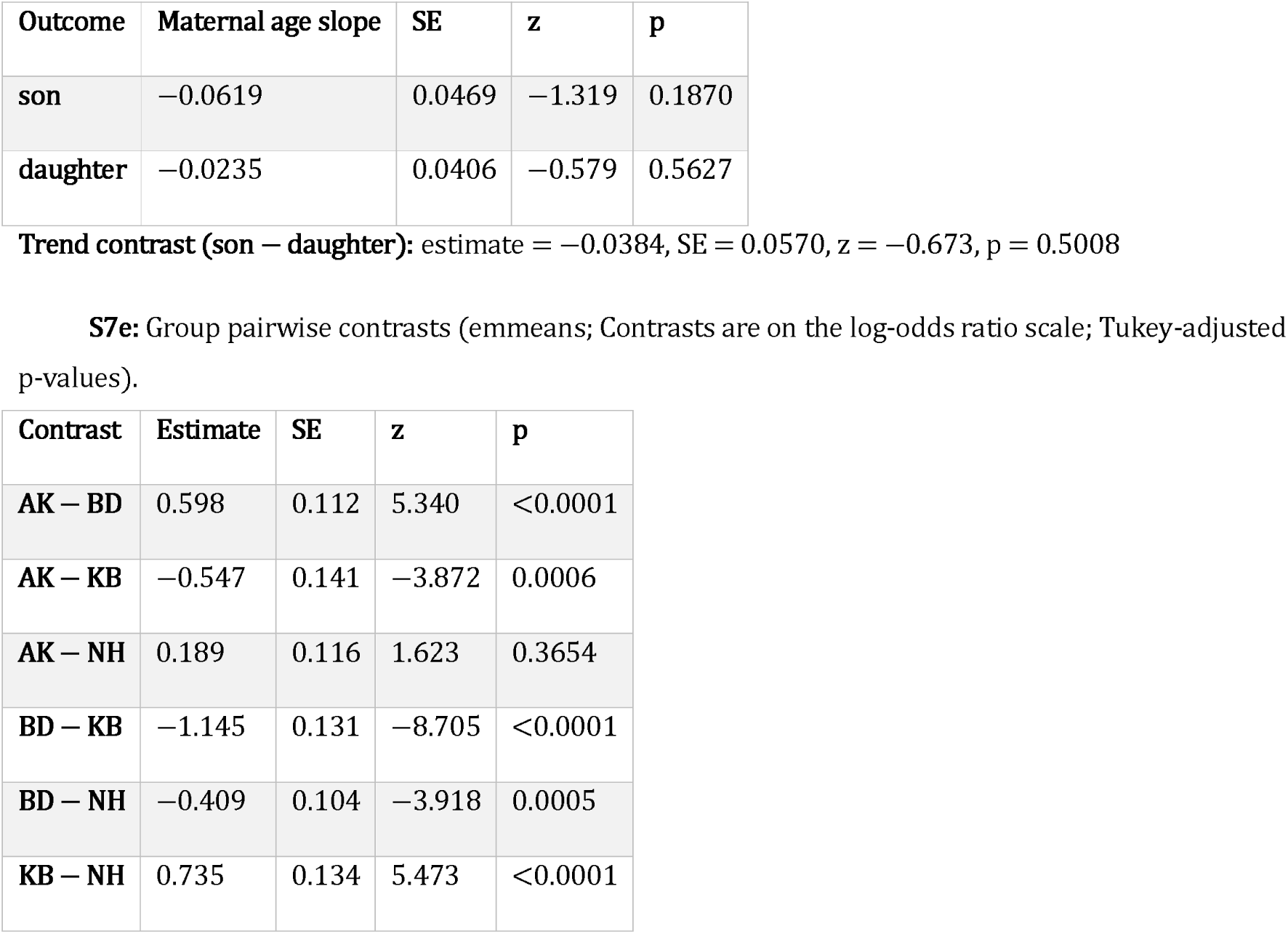
Grooming model (betabinomial GLMM; response = cbind(successes, trials-successes)). Table S7a: Fixed predictor outcomes.

**Table S8:**
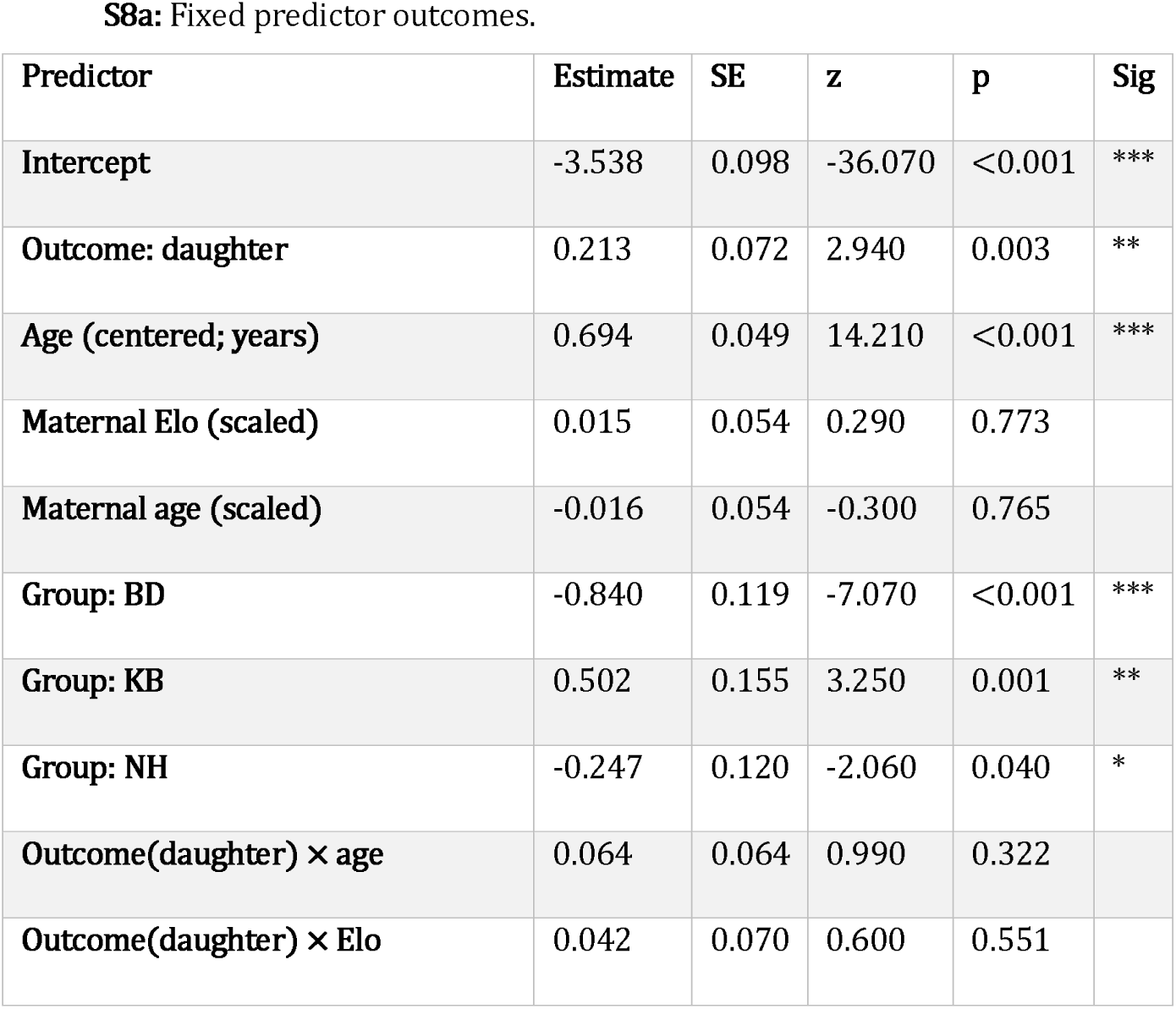
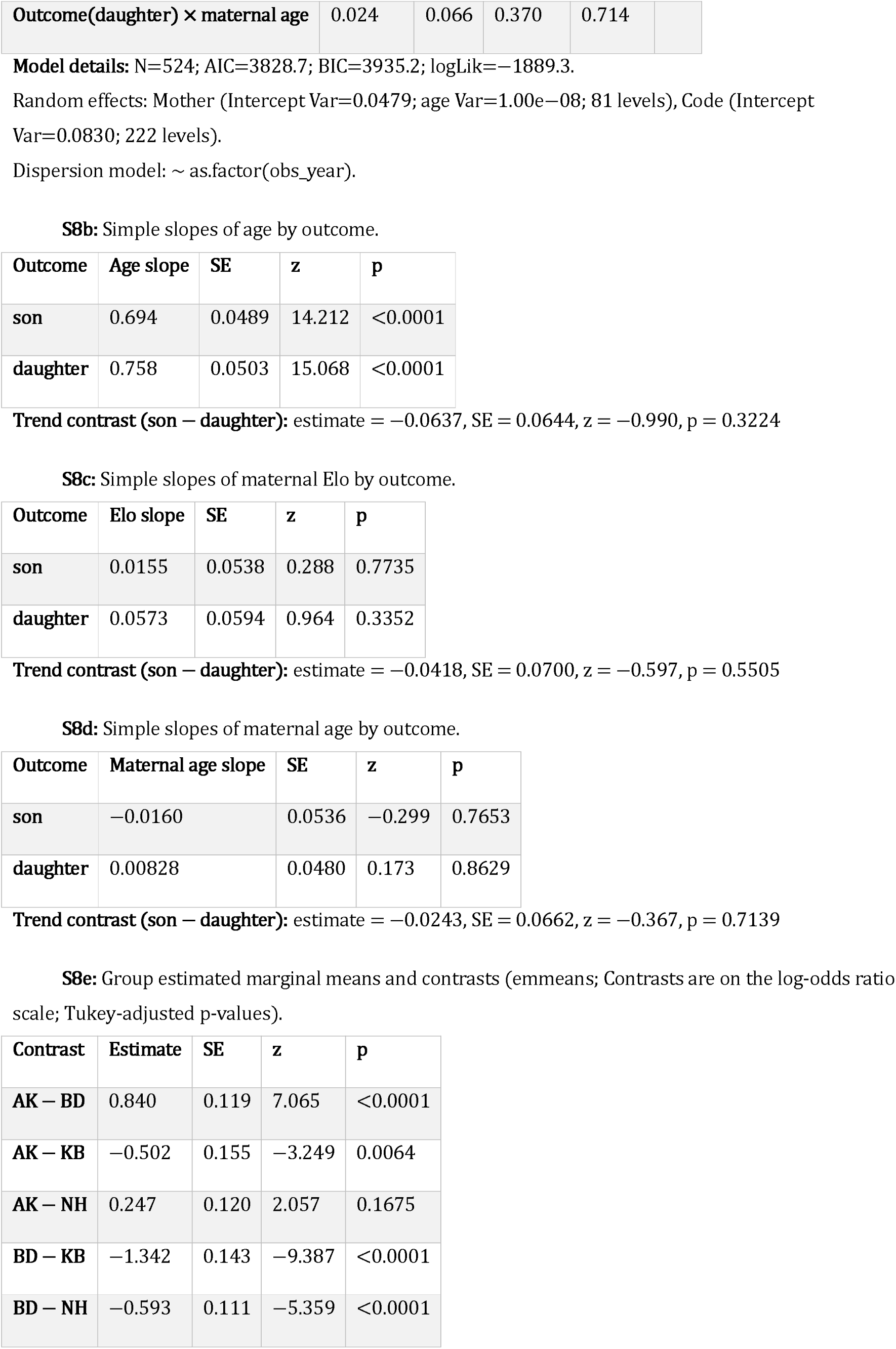
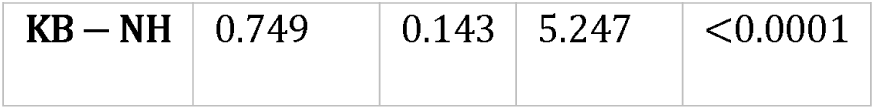
Conflict model (betabinomial GLMM; response = cbind(successes, trials−successes); trials ≠ 0). Table S8a: Fixed predictor outcomes.

**Table S9:**
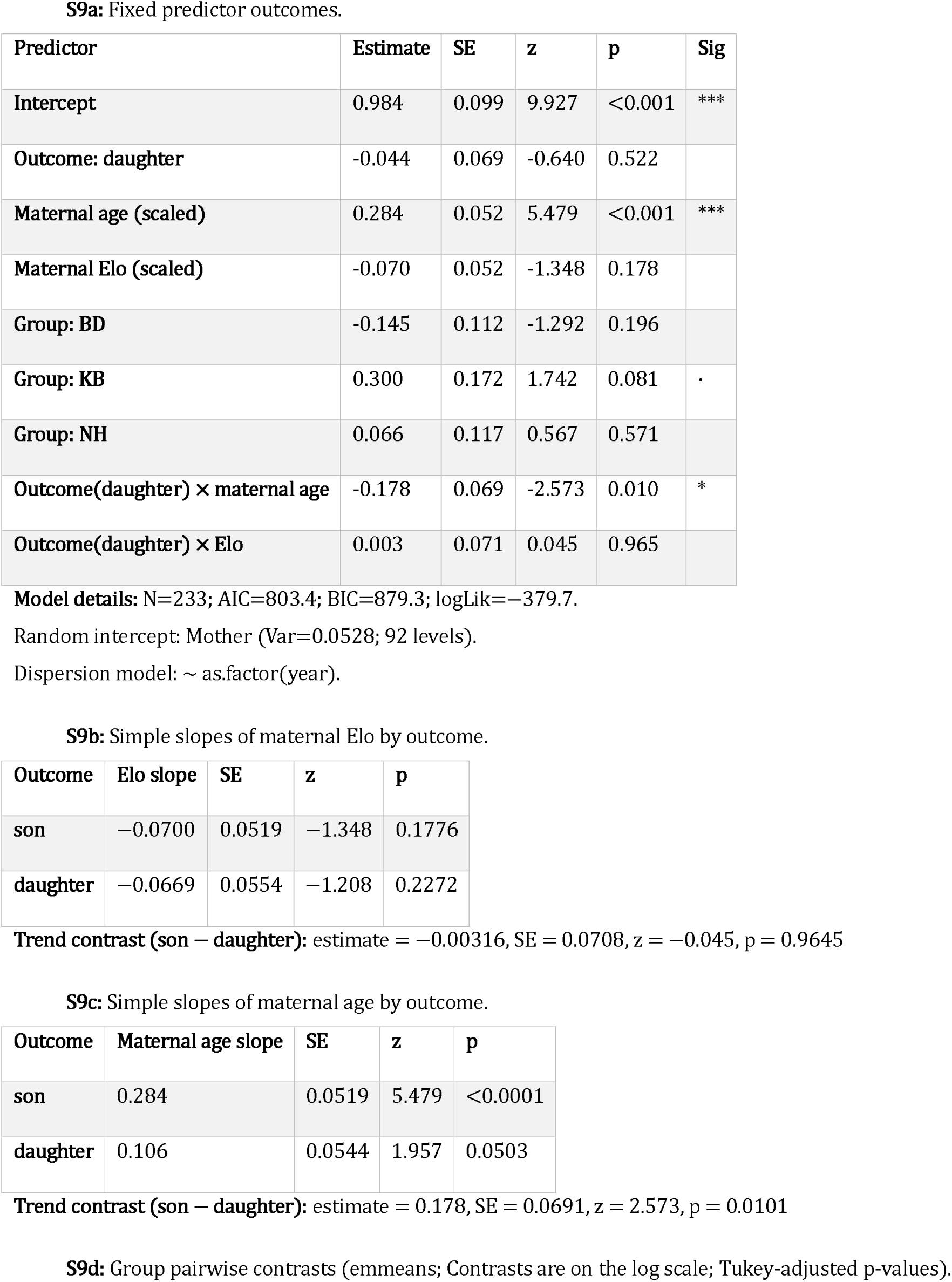
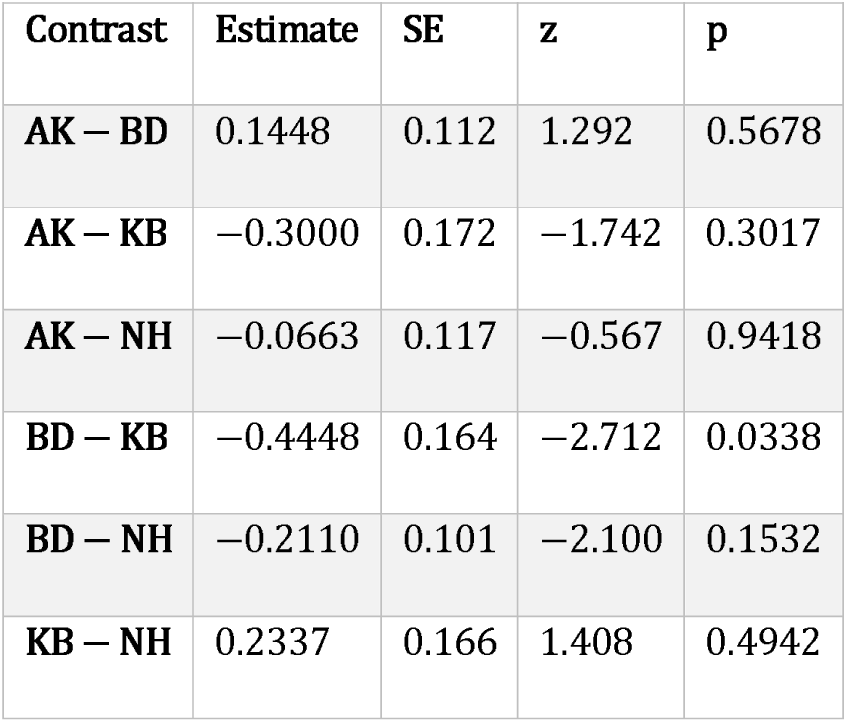
Nearest-adult distance model (Gamma GLMM; response = mean closest adult distance; log link).

**Figure S1:**
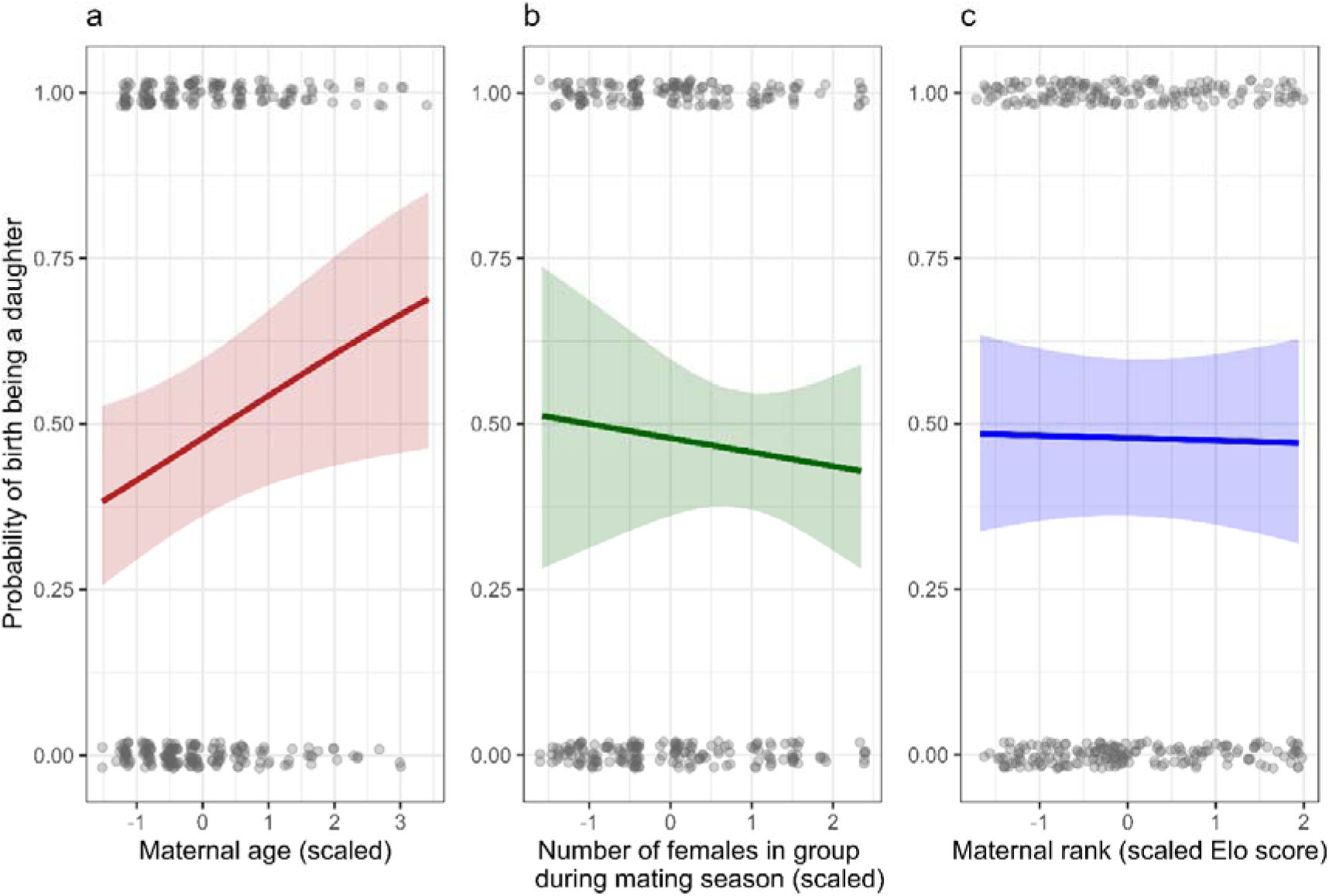
Predictors of offspring sex ratio at birth. Model-predicted probability that a birth was a daughter (1 = daughter, 0 = son) plotted against maternal and social predictors. Points show observed birth outcomes (0/1, jittered; raw data points); solid lines show model predictions and shaded ribbons indicate 95% CIs. (a) Maternal age at birth (scaled): probability of producing a daughter increased with maternal age. (b) Number of females present in the group during the mating season (scaled): no evidence that local female abundance predicted offspring sex. (c) Maternal dominance rank (scaled Elo score): no evidence that maternal rank predicted offspring sex at birth.

**Figure S2:**
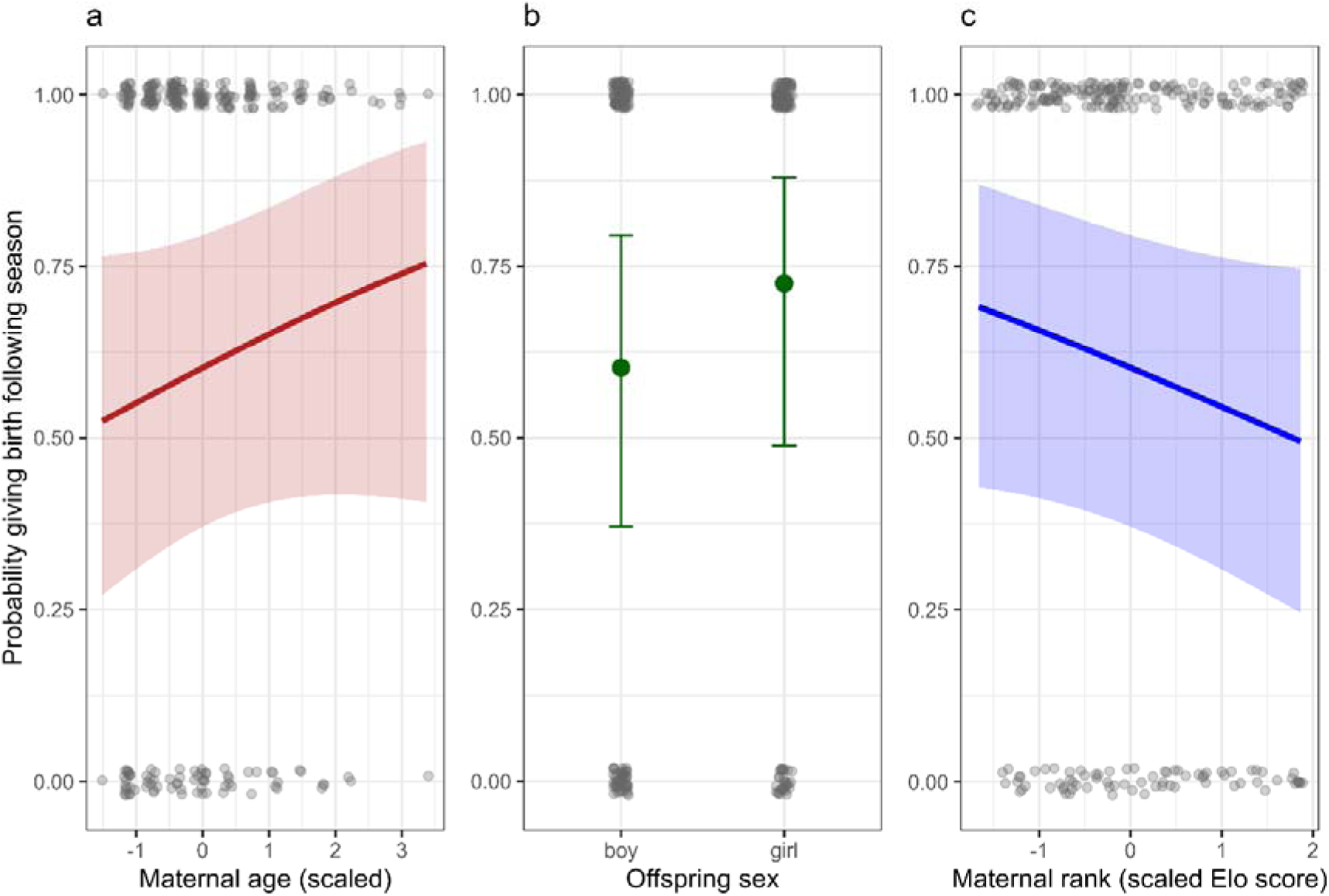
Predictors of reproductive pace. Model-predicted probability that a mother gave birth in the breeding season following her previous birth (1/0), shown across maternal characteristics and offspring sex. Points show observed reproductive outcomes (0/1, jittered; raw data points); lines and shaded ribbons show model predictions ± 95% CI, and points with error bars in (b) indicate predicted means ± 95% CI. (a) Maternal age (scaled): no evidence that maternal age predicted giving birth the following season. (b) Offspring sex (son/daughter): mothers were marginally more likely to reproduce again after giving birth to daughters than to sons. (c) Maternal dominance rank (scaled Elo score): no evidence that maternal rank predicted giving birth the following season.

**Figure S3:**
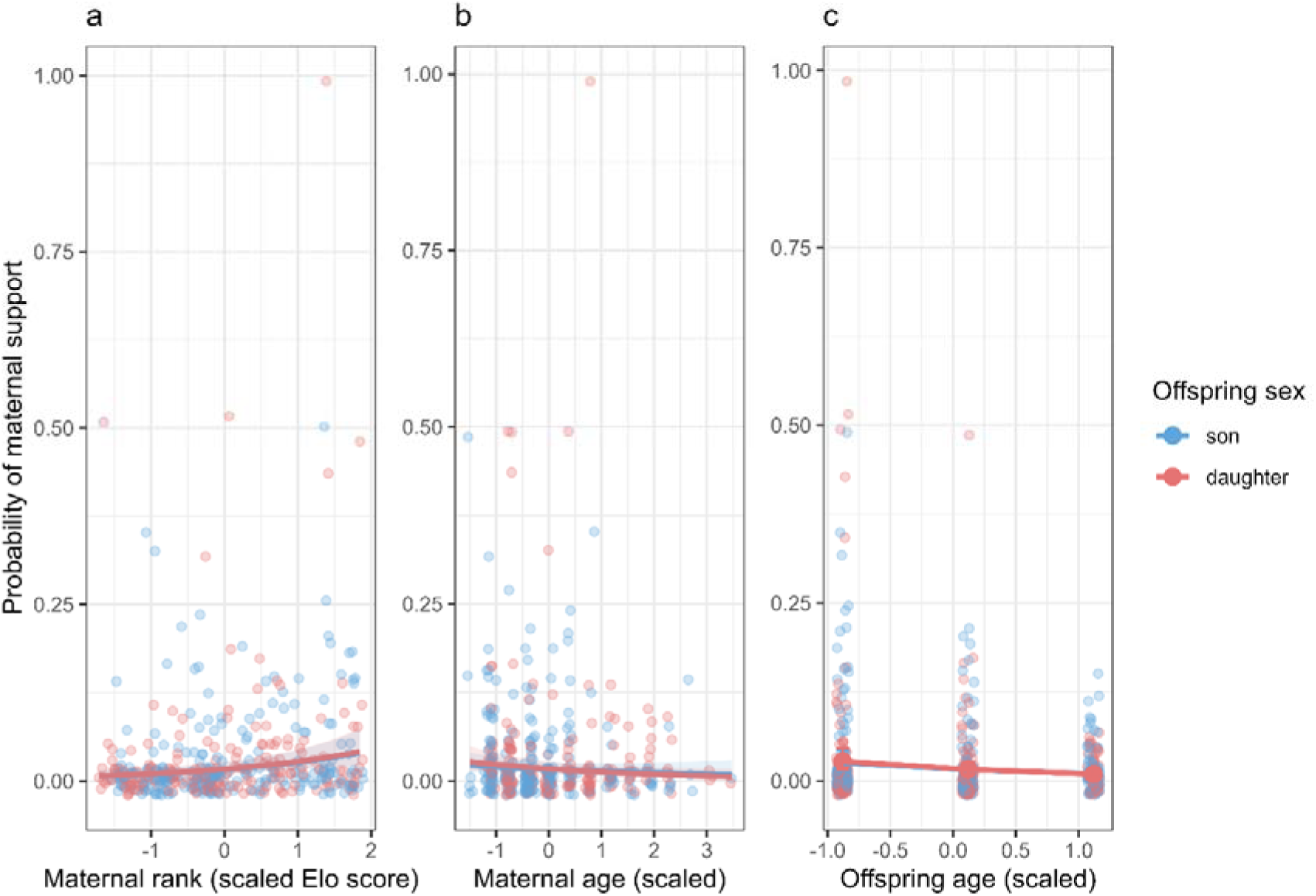
Maternal coalitionary support toward offspring across maternal rank, maternal age, and offspring age. Model-predicted probability that a mother supported her offspring in a conflict (1/0) plotted against maternal rank, maternal age, and offspring age, shown separately for sons (blue) and daughters (red). Points represent individual conflict observations (jittered; raw data points); lines show model predictions with shaded 95% confidence intervals. (a) Maternal dominance rank (scaled Elo score): probability of maternal support increased with maternal rank. (b) Maternal age (scaled): probability of maternal support decreased with maternal age. (c) Offspring age (scaled): probability of maternal support decreased with offspring age.

